# A cyclic dipeptide for salinity stress alleviation and the trophic flexibility of an endophyte reveal niches in salt marsh plant-microbe interactions

**DOI:** 10.1101/2023.12.12.569982

**Authors:** Shih-Hsun Walter Hung, Pin-Hsien Yeh, Tsai-Ching Huang, Shao-Yu Huang, I-Chen Wu, Chia-Ho Liu, Yu-Hsi Lin, Pei-Ru Chien, Fan-Chen Huang, Ying-Ning Ho, Chih-Horng Kuo, Hau-Hsuan Hwang, En-Pei Isabel Chiang, Chieh-Chen Huang

**Affiliations:** Department of Life Sciences, National Chung Hsing University, Taichung 402202, Taiwan; Institute of Plant and Microbial Biology, Academia Sinica, Taipei 115201, Taiwan; Advanced Plant and Food Crop Biotechnology Center, National Chung Hsing University, Taichung 402202, Taiwan; Institute of Marine Biology, College of Life Science, National Taiwan Ocean University, Keelung 202301, Taiwan; Center of Excellence for the Oceans, National Taiwan Ocean University, Keelung 202301, Taiwan; Biotechnology Center, National Chung Hsing University, Taichung 402202, Taiwan; Innovation and Development Center of Sustainable Agriculture, National Chung Hsing University, Taichung 402202, Taiwan; Department of Food Science and Biotechnology, National Chung Hsing University, Taichung 402202, Taiwan

**Keywords:** endophyte, cyclic dipeptide, molecular plant-microbe interactions (MPMI), symbiosis

## Abstract

In response to climate change, the nature of endophytes and their applications in sustainable agriculture has attracted the attention of academia and agro-industries. We focused on the endophytic halophiles of the endangered Taiwanese salt marsh plant, *Bolboschoenus planiculmis*, and evaluated the functions of the isolates through *in planta* salinity stress alleviation assay using *Arabidopsis*. An endophytic strain *Priestia megaterium* BP01R2 that could promote plant growth and salinity tolerance was further characterised through multi-omics approaches. The transcriptomics results suggested that BP01R2 could function by tuning hormone signal transduction, energy-producing metabolism, multiple stress responses, etc. In addition, a cyclodipeptide, cyclo(L-Ala-Gly), identified by metabolomics analysis was later confirmed to contribute to salinity stress alleviation in stressed plants by exogenous supplementation. Here we provide a new perspective on host-microbe interactions in the wetland biome based on the multi-omics investigation and mixotrophic character of BP01R2. This study revealed a biostimulant-based plant-endophyte symbiosis with potential application in sustainable agriculture and facilitated our understanding of those enigmatic cross-kingdom relationships.

## Introduction

Climate change and projected world population growth threaten our food supply systems (1,2). To ensure adequate food security and sustainability, it’s crucial to develop new technologies for growing crops efficiently in extreme environments (3). Apart from traditional breeding and precision genetic technologies, plant growth-promoting rhizobacteria (PGPRs) and endophytes have also been implemented to enhance crop nutrient and water absorbance, hormone signal transduction, stress adaptation, etc. (4,5). Among them, endophytes that live inside plants (endosphere) are attracting attention due to their *in planta* colonisation properties and specific effects on host plants for growth promotion and tolerance to both abiotic and biotic stresses (6–9). Various endophytes isolated from diverse origins were demonstrated to exhibit positive effects by producing bioactive metabolites known as biostimulants (10–15).

Endophytes isolated from saline habitats, such as salt marshes or alkaline lakes, may be promising candidates for endophyte-assisted agriculture in saline or overfertilized fields. For the isolation of such endophytic halophiles, plants growing in conditions with continuously high salinity and periodic hypoxia are suitable sources (16–18). Salt marsh plants often face multiple stresses, but many wetland-related studies have been limited to microbe-assisted phytoremediation (19–22). Therefore, understanding the strategies used by these unique plant-associated microbes living in stressful environments through symbiosis mechanisms may provide novel knowledge and tools for future applications.

In previous studies, we isolated endophytic strains from various natural habitats and used them to construct synthetic plant-endophyte symbiosis systems for plant cultivation under biotic or abiotic stress conditions (5,23–26). On the other hand, some specific metabolites, such as pyrroloquinoline quinone (PQQ) and cyclodipeptides (CDPs), have been identified as potential biostimulants (23,27–30), while some CDPs were also reported as agricultural agents that could protect crops against biotic and abiotic stresses (31–35). Although some of them had been patented for applications more than three decades ago (36), the molecular bases of the role of cyclodipeptides in moderating plant stresses and in plant‒microbe interactions remain to be elucidated.

Here we isolated and characterised halotolerant and PGP endophytes from endangered salt marsh plants in Taiwan. The complete genome and metabolome of *Priestia megaterium* BP01R2 (previously known as BP-R2), plus its *in planta* regulations on host transcriptome, were further investigated to reveal the molecular mechanism among such symbiotic relationships. In the meantime, a cyclodipeptide cyclo(L-Ala-Gly) was identified as a novel CDP biostimulant for plant salinity stress alleviation. Overall, this study facilitates our understanding of the fundamental science of the enigmatic cross-kingdom host-microbe interactions and unveils promising endophytic biostimulants for sustainable agriculture.

## Materials and Methods

### Plant materials and sampling

The plant material (*Bolboschoenus planiculmis*) for endophyte isolation was collected from a tidal marsh in Taichung, Taiwan (Fig. 1A). The seeds of *Arabidopsis thaliana* ecotype Columbia (Col-0) were retrieved from Chieh-Chen Huang Lab’s stock.

**Fig. 1.**
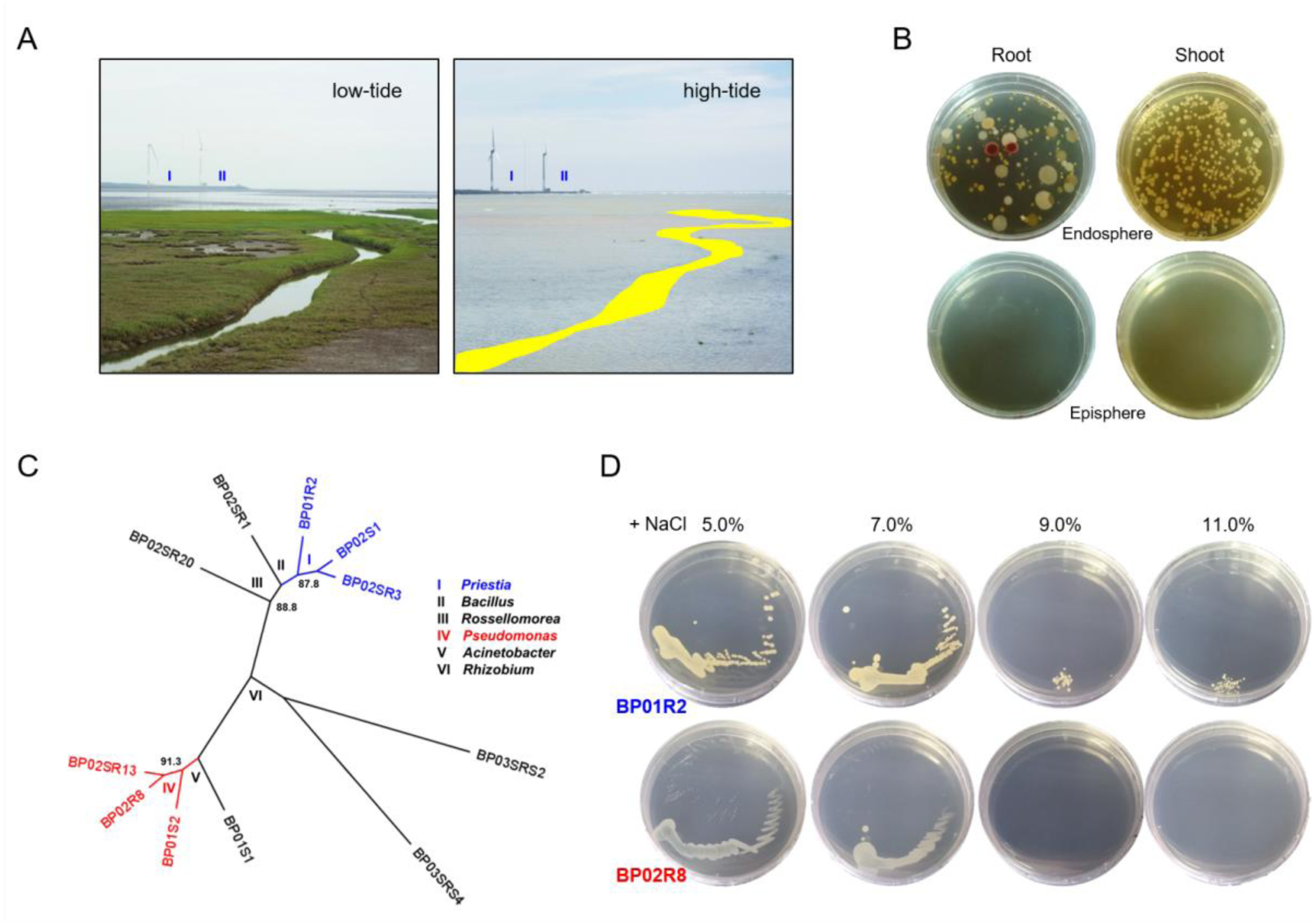
Typical habitat and bacterial endophyte identification of *Bolboschoenus planiculmis*. **A** Typical niche, tidal marsh in Taichung, Taiwan at low-(left) and high-tide (right). I and II show two corresponding locations in two different photographs. The pseudo-colour marks indicate a channel visible in the low-tide. **B** Endophyte microbiome. Bacterial endophytes isolated from different plant tissues (Endosphere). Control check of last rinsing solution used for surface sterilising (Episphere). **C** Unrooted 16S rRNA gene-based maximum likelihood phylogeny of eleven bacterial candidates. All internal nodes received > 85% bootstrap support based on 1,000 re-samplings. Roman numerals indicate that the groups belong to different genera. The groups containing isolates BP01R2 and BP02R8 focused on in this work are shown in blue and red, respectively. **D** *in vitro* salinity stress tolerance test of endophytes.

### Endophyte isolation, characterisation and 16S rRNA gene-based phylogeny

Endophytic bacteria isolation and maintenance were performed as in our previous works (23,26). In brief, the endosphere fluids of surface-sterilised plant tissues were used for endophyte isolation and incubation using Luria-Bertani (LB) agar or broth at 37°C for 24-48 hours. The isolates were first picked based on their different morphologies, and conducted a preliminary taxonomic check by 16S rRNA gene-based phylogeny. The bacterial IAA production was examined as described by Hwang *et al.* (26). The 16S rRNA genes were amplified using the primer set E8F/U1510R as previously described (5,26). Multiple sequence alignments of 16S rRNA genes were performed using MUSCLE v3.8.31 (37). Maximum likelihood phylogenies were inferred using PhyML v3.3 (38) and visualised using FigTree v1.4.4. PHYLIP v3.697 (39) was used for bootstrap analysis.

### Plant-endophyte symbiosis and biostimulants assays

Unless otherwise stated, *Arabidopsis* was grown according to previously described methods (26,40). In short, seeds were germinated on Murashige and Skoog (MS) plates with NaCl (ranging from 0 mM to 170 mM) under a 10/14 hour day/night cycle at 25°C. Endophytes were incubated and inoculated to the plants as described in our previous work (26). Briefly, strains BP01R2 and BP02R8 were adjusted on OD_600_ = 0.8, ≈ 3.5 x 10^7^ and 9.4 x 10^8^ CFU/mL, respectively, then diluted into OD_600_ = 0.4 and 0.04 as the relatively low (LCI) and high concentration inocula (HCI) for the symbiosis assays. The biostimulant cyclo(L-Ala-Gly) (SS-2476, Combi-Blocks; San Diego, California, United States) was purchased from UNI-ONWARD Corp (New Taipei City, Taiwan) and was prepared for the plant experiments.

### Plant total RNA extraction and transcriptomic analysis

*Arabidopsis* seedlings grown on MS plates supplemented with 100 mM NaCl were harvested 15 days after inoculation (DAI). Samples were homogenised with mortar and pestle using liquid nitrogen; the total RNA was extracted from the mixtures of five seedlings of each treatment using TRIzol reagent. The library preparation and RNA sequencing were performed following Welgene Biotech’s in-house pipeline as in our previous works on plant transcriptomes (41,42); all kits were used according to the manufacturer’s instructions, and all bioinformatics tools were used with the default settings. The libraries construction was carried out with Agilent’s SureSelect Strand-Specific RNA Library Preparation Kit followed by size selection with AMPure XP beads (Beckman Coulter; Chaska, Minnesota, United States) and 75 bp single-end sequencing was performed on the Illumina Solexa platform. Illumina’s program bcl2fastq v2.20 was used for basecalling and low-quality reads were trimmed off based on Q20 accuracy. The resultant sequence was mapped to the TAIR10 genome (43). The transcript per million (TPM) method was used for normalisation, and genes with TPM < 0.3 were excluded (44). Genes with 2.0-fold TPM differences and a probability of at least 0.95 were defined as differentially expressed genes (DEGs). StringTie v2.1.4 and DEseq2 v1.28.1 were used for genome bias detection/correction (45,46). Gene set enrichment analysis (GSEA) (47) was carried out, and the DEGs were accessed to Gene Ontology (GO) (48) and Kyoto Encyclopedia of Genes and Genomes (KEGG) (49) database for the metabolic pathways prediction.

### Bacterial metabolomics

The strain BP01R2 was cultured on LB agar plates with or without NaCl, as mentioned above. Five agar plates were collected for each sample, and metabolites were extracted with ethyl acetate two times. The extracts were re-dissolved in methanol and adjusted to 10 mg/mL. All samples were analysed using the linear ion trap mass spectrometer system (LTQ XL, Thermo Fisher Scientific; San Jose, California, United States) with direct injection at positive ion mode. The mass range was from m/z 100-1500. MS raw data files were converted to mzXML format using MSConvert (50). Metabolite peak detection was performed using MZmine version 2.53 (51), and the pre-processed data was conducted by R program (version 4.0.3) (52). The pre-processed MS feature table and unknown metabolites were searched and annotated against referenced metabolites of the AntiMarin database (53) by exact molecular mass to identify the molecular formula and for annotation of discriminating features.

### Bacterial genome sequencing, assembly and annotation

The procedures for genome sequencing and analysis were based on those described in our previous work on bacterial genomes (29,54). All kits were the same as previously described and were used according to the manufacturer’s protocols, and all bioinformatics tools were used with the default settings unless stated otherwise. Briefly, the total DNA of strain BP01R2 was extracted and then sequenced via Illumina NovaSeq 6000 2 × 150 bp paired-end and Oxford Nanopore Technologies (ONT) MinION platforms. Further filtering was conducted to remove ONT reads shorter than 12,000 bp. A hybrid *de novo* assembly was produced using Unicycler v0.4.9-beta (55). For validation, the Illumina and ONT raw reads were mapped to the assembly using BWA v0.7.17 (56) and Minimap2 v2.15 (57), respectively. The results were programmatically checked with SAMtools v1.9 (58) and manually inspected using IGV v2.11.1 (59). For assembly completeness evaluation, benchmarking universal single-copy orthologous (BUSCO v5.1.2) analysis, referenced to the Bacillales dataset, was executed on gVolante (60,61).

The finalized assembly was submitted to the National Center for Biotechnology Information (NCBI) and annotated using the Prokaryotic Genome Annotation Pipeline (PGAP) (62). KofamKOALA (63) was used to examine putative metabolic pathways.

### Bacterial comparative genomics

All BioProject and BioSample records of *P. megaterium* accessible on NCBI as of Feb. 28, 2022 were retrieved for comparative analysis with BP01R2 (Table S1). FastANI v1.1 (64) was used for whole-genome comparison to calculate the proportion of genomic segments mapped and the average nucleotide identity (ANI). Homologous gene clusters were identified using OrthoMCL (65) for gene content analysis. The genes associated with plant growth-promoting (PGP) traits and rhizosphere competence were identified according to previous works on *P. megaterium* (66) and bacterial endophytes (67); genes associated with salinity stress (68–70) and autotrophs (71–74) were also identified. Multiple sequence alignments of homologous genes, maximum likelihood phylogenies and bootstrap analysis were conducted based on the methods described for 16S rRNA gene phylogeny.

### Bacterial tropism analysis

Anaerobic cultivation was performed according to the instructions of the Leibniz Institute DSMZ (75). The Hungate anaerobic culture tube and Coy anaerobic chamber (Coy Laboratory Products, United States) were used in this experiment. BP01R2 was pre-cultivated 24 hours in M9 minimal medium (248510; Becton Dickinson, United States), and then washed three times using the same method before use as an inoculant. The inoculant density was set at OD_600_ = 0.01; the cultivation volume was 5 mL, and cultivation was carried out with shaking at 120 rpm and 37℃. M9 minimal medium supplemented with 2 mM MgSO_4_ (131-00405; FUJIFILM Wako, Japan), 0.1 mM CaCl_2_ (21075; Sigma-Aldrich, United States), 10 mM NaNO_3_ (195-02545; FUJIFILM Wako, Japan) and 0.01% L-tryptophan (A10230; Alfa Aesar, United States) was prepared as a general incubation medium. Extra glucose as an organic carbon resource was tested at concentrations of 0.4%, 0.2%, 0.04% and 0%; the tube headspace gas was composed of either sterile air or 90% H_2_ plus 10% CO_2_ (N_2_). The growth curves were measured at OD_595_ using the PARADIG detection platform (Beckman, United States) or Sunrise absorbance microplate reader (Tecan, Austria).

### Statistics

The Shapiro-Wilk and Levene’s tests were used to check the normal distribution of variables and homoscedasticity. A parametric one-way ANOVA analysis with Tukey’s post hoc HSD test or non-parametric Kruskal-Wallis test with Dunn’s post hoc test was then applied to evaluate statistical significance. The data analysis was performed using the Real Statistics Resource Pack v7.7.1 (76). For all experiments, at least five independent biological replicates were tested and data are shown as mean±SEM unless otherwise stated. For all data points, *P ≤ 0.05, **P ≤ 0.01 and ***P ≤ 0.005, indicated significant differences between samples; otherwise, not significant.

## Results

### Halotolerant bacterial endophytes isolation

Totally 128 and 519 colony-forming units (CFUs) per 100 μL were found in the endosphere of roots and shoots, respectively; however, the microbiome composition in shoots was less diverse (Fig. 1B). Among the picked isolates, 11 endophytes consisted of six genera: *Priestia*, *Pseudomonas*, *Rhizobium*, *Bacillus*, *Acinetobacter*, and *Rossellomorea* (ordered from high to low enrichment) (Fig. 1C). A *Priestia* sp. strain (designated as strain BP01R2) and a *Pseudomonas* sp. strain (designated as strain BP02R8) were chosen for the following experiments due to their enriched distribution and cases of beneficial endophytes affiliated to these genera (66,77,78). For the halotolerant test, both strains tolerate 7.0% NaCl supplementation, and BP01R2 can even survive under an extremely high concentration of 11% NaCl, which was therefore identified as a halophile (Fig. 1D).

### Plant-endophyte symbiosis

The symbiosis assay was first tested without salinity stress. Both BP01R2 and BP02R8 presented plant-growth-promotion characters either been prepared as low (LCI) or high concentration inocula (HCI) (Fig. 2A). At 15 days after inoculation (DAI), both strains improved the plant main root length (Fig. 2B) and fresh weight accumulation (Fig. 2C). BP01R2 and BP02R8 induced the formation of plant lateral roots under HCI (Fig. 2A, D). BP01R2 also caused more root hairs under LCI, and the statistical significance in plant main root elongation was found for both strains under HCI. These results echoed the previous observation of strains on plant growth promotion (26) and highlighted their potential benefits for rhizosphere competence.

**Fig. 2.**
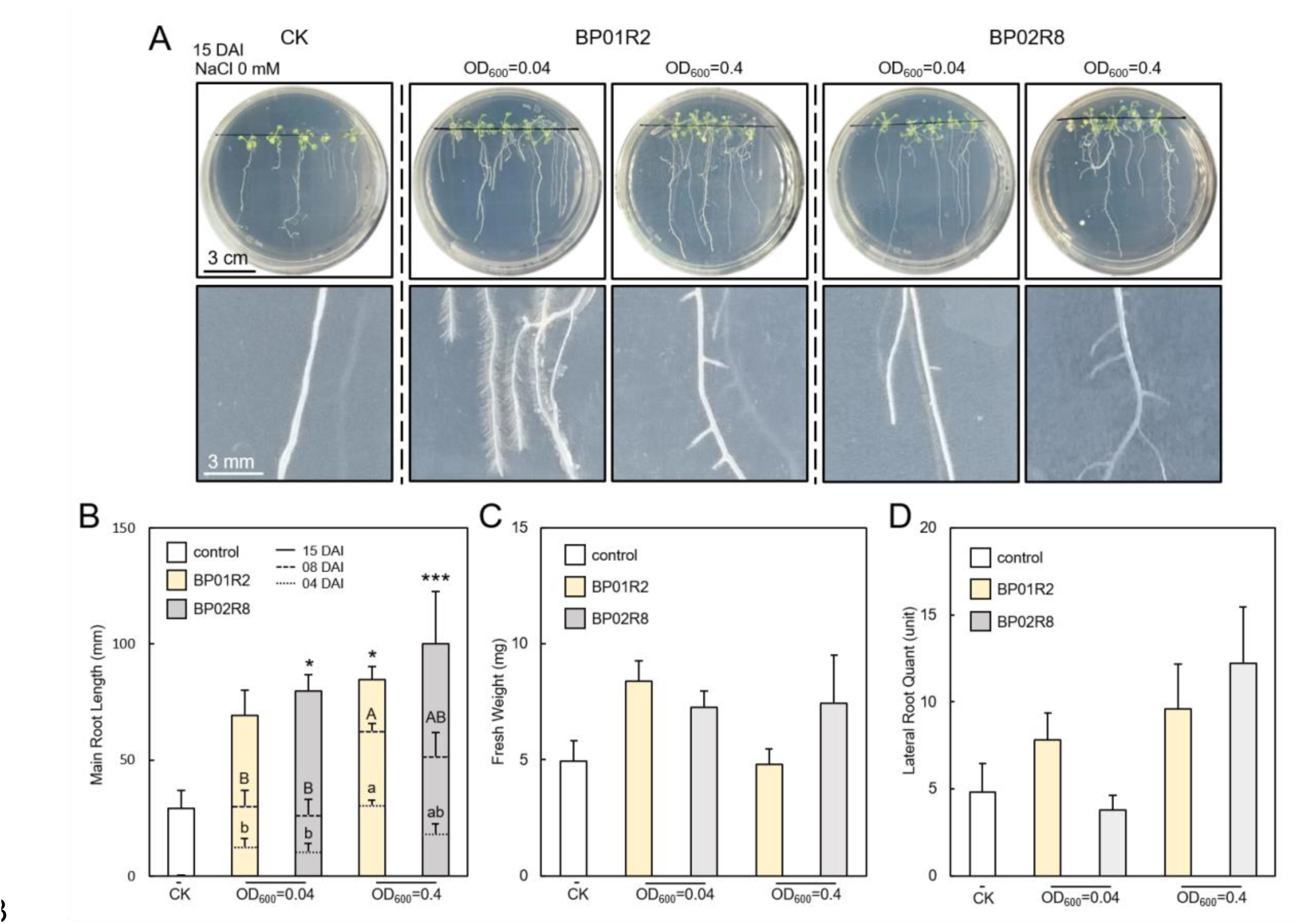
Endophytes promote host plant growth. **A** Plant growth promotion differs with inoculation concentration and type of endophyte. The upper five panels illustrate the plant growth patterns under treatment with different inocula, and the lower five panels provide a close-up view of the roots. CK: control check; DAI: days after inoculation. **B** The main root length, **C** fresh weight and **D** lateral root number of different inoculum-treated plants. Data are presented as mean±SEM. N=5. Different letters show the significant difference between samples (p<0.05); 04 DAI are uncapitalised; 08 DAI are capitalised. For 15 DAI, *p<0.05 ***p<0.005.

### Endophytic biostimulants for *in planta* salinity stress alleviation

The seedlings of *A. thaliana* were grown in MS medium supplemented with 85 mM and 170 mM NaCl for 15 days to mimic salinity stress. BP01R2 and BP02R8 benefited the plant growth on shoots, main root and fresh weight, just like in stress-free conditions (Fig. 3). Phenotypes of chlorotic and adaxial-curled leaves, fewer lateral roots and shorter main root length observed in the control plant (CK) under 170 mM NaCl stress were mitigated through endophyte inoculations (Fig. 3A). For instance, the decreasing ratio of the main root length of CK, BP01R2 and BP02R8 were 56.2%, 15.2% and 17.9%, respectively. On the other hand, both endophytes improved the main root length, and BP01R2 further alleviated the decrease of the main root length under 170 mM NaCl salinity stress (Figs 2AB, 3AE). Also, two endophytes significantly increased the plant’s fresh weight under 85 mM and/or 170 mM NaCl stress (Fig. 3C, D).

**Fig. 3.**
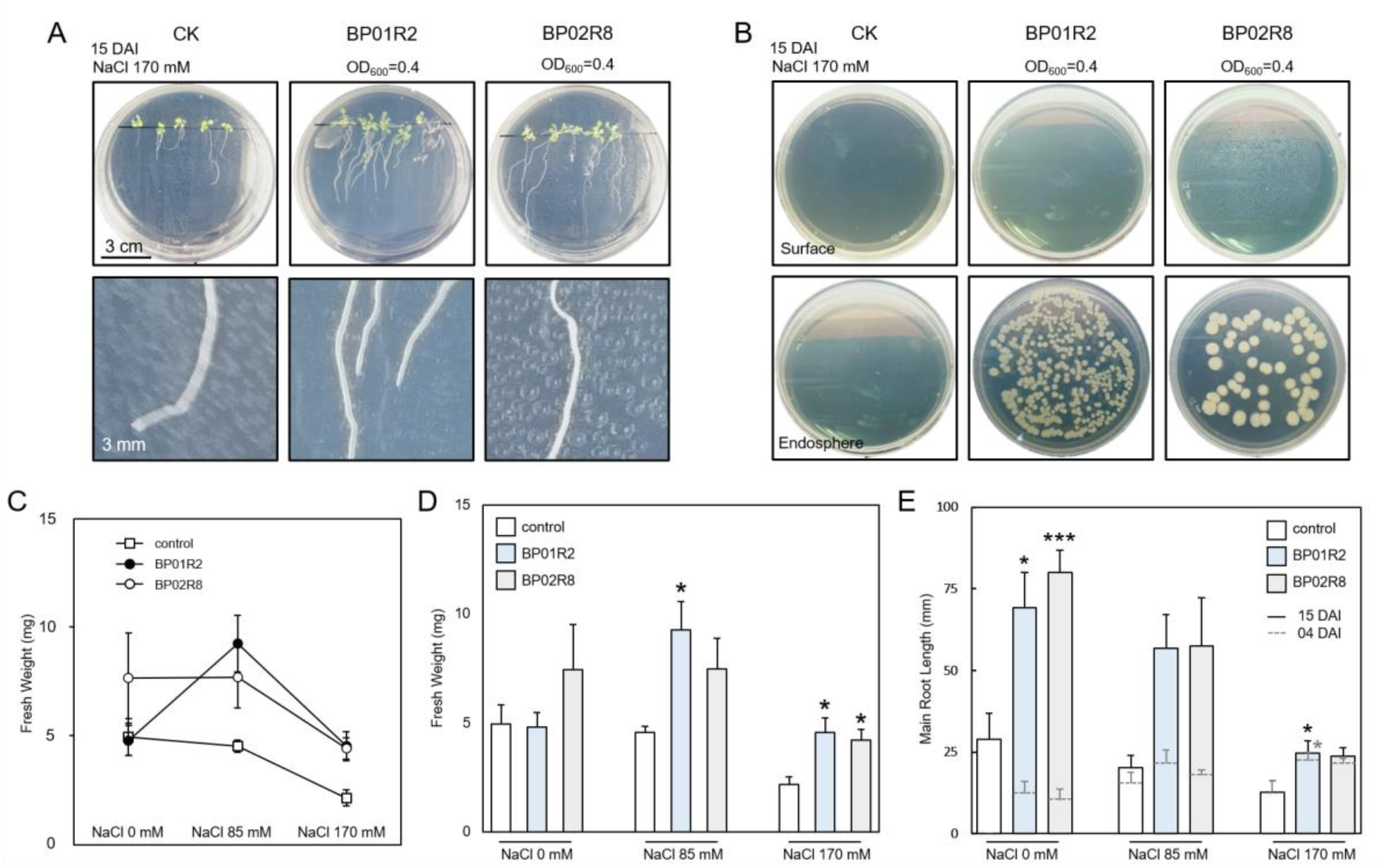
Endophytes alleviate salinity stress in the host plant. **A** Plant growth differs from endophytic inoculation under salinity stress. The upper three panels illustrate the plant growth patterns with different inocula under salinity stress, and the lower three panels provide a close-up of the roots. CK: control check; DAI: days after inoculation. **B** Endophytes re-isolation. **C D** The fresh weight (15 DAI) and **E** main root length (04 and 15 DAI) of plants treated with different inocula under salinity stress. Data are presented as mean±SEM. N=5 *p<0.05 ***p<0.005.

For the transcriptomic analysis, a total of 30,956,567 (∼2.3 Gb), 34,330,286 (∼2.6 Gb), 31,539,938 (∼2.4 Gb) and 32,554,093 (∼2.4 Gb) trimmed reads were generated for CK, BP01R2, CK_NaCl and BP01R2_NaCl, respectively. Strikingly, the expression pattern of the CK was intermediate to the samples that suffered from salinity stress, and the BP01R2_NaCl showed a higher similarity to the CK instead of the CK_NaCl, which may result from the alleviation contributed by the symbiosed endophyte as seen in the phenotypic data (Fig. 4B). When the plant-BP01R2 symbiosis occurred, totalling 658 DEGs within 21,028 genes were found (Fig. S1A), and the gene ontology (GO) analysis showed the biological processes of plant epidermal and root cell differentiation and root morphogenesis; the cellular component of chromosome, cytoskeleton, transferase, and cytoplasmic vesicle; the molecular function of oxygen binding, ATPase activity, helicase activity, and ligase activity was enrich-regulated (Figs S2, S3). Consistent with the GO enrichment result, several PGP-related pathways (*e.g.*, glycolysis and gluconeogenesis, oxidative phosphorylation, plant hormone signal transduction, etc.) were also shown to be enrich-regulated within the KEGG analysis result (Figs S4-S7). These enriched gene regulations well explained the phenotypes observed in the symbiosis assays (*i.e.*, the growth improvement traits such as main root elongation, lateral root induction, fresh weight accumulation, etc.) (Figs 2, 3A,C-E).

**Fig. 4.**
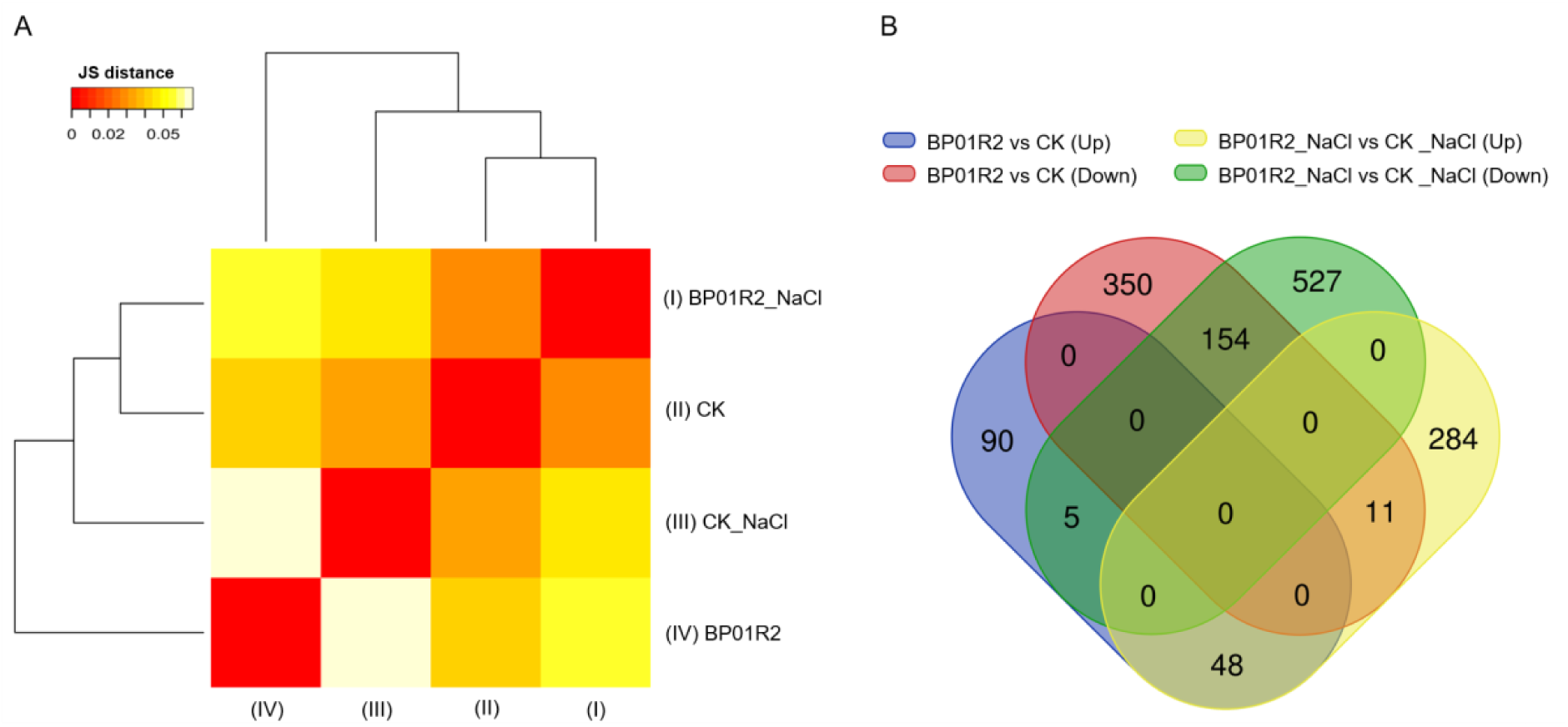
Transcriptomic analysis. **A** Heatmap of Jensen-Shannon (JS) divergence between transcriptomics data for all genes. **B** Venn diagram of DEGs comparison between samples from with or without salinity stress.

When the symbiosis occurred under salinity stress, a total of 1029 DEGs were identified among 20,937 genes (Fig. S1B). The cell death, hypersensitive response, jasmonic acid biosynthesis and response, ethylene response, MAPK signalling, and glutathione metabolism, among others, were up-regulated (Figs S8-S13) and usually exhibited opposite results compared to the CK_NaCl and CK comparison (Figs S14-S17). Meanwhile, the plant hormone signal transduction and ribosome-related pathways were consistently up-regulated, like the expression pattern for symbiosis without the salinity stress. Furthermore, the simultaneously up- (co-up-DEGs) and down-regulated DEGs (co-down-DEGs) between samples with and without salinity stress were also examined. A total of 202 co-DEGs were universally up- or down- regulated, regardless of salinity stress or not (Fig. 4B). Among them, the 48 co-up-DEGs were found to be involved in plant-type cell wall organization or biogenesis, xyloglucan:xyloglucosyl transferase activity, peroxidase activity, oxidoreductase activity, antioxidant activity, etc. (Table S2); the 154 co-down-DEGs were enriched in response to hypoxia, jasmonic acid, salicylic acid, abscisic acid, oxidoreductase, oxidative stress, and related metabolism (Table S3). In summary, the transcriptome data are consistent with the phenotypes and explain at the molecular level how BP01R2 symbiosis alleviates *in planta* salinity stress and promotes the host plant growth.

### Cyclo(L-Ala-Gly) improves plant growth and alleviates *in planta* salinity stress

To investigate which metabolites could contribute to hosts in salinity stress alleviation, we used the linear ion trap mass spectrometer system to analyse the metabolomics of BP01R2 under NaCl-present and -absent incubating conditions. Totally 191 compounds were identified specifically under the NaCl-present condition, and 249 compounds were found to be shared by NaCl-present and -absent conditions (Fig. 5A, Tables S4, S5). Among the profiles, 18 cyclic dipeptides, also known as 2,5-diketopiperazines and cyclo dipeptides (CDPs), were identified. These CDPs could be classified into proline-(includes hydroxyproline) and non-proline-based groups and nine were observed only under the NaCl-present condition (Table S6). CDPs have been reported as biostimulants against biotic and abiotic stresses in plants (31–35), and many proline- and hydroxyproline-based CDPs were patented in inducing resistance in plants against abiotic stresses (36). However, our understanding of those non-proline-based CDPs and their roles in MPMI remains limited. Therefore, we focused on the only non-proline-based CDP, cyclo(L-Ala-Gly), produced by BP01R2 under NaCl-present conditions for the follow-up biostimulants assays. After three weeks of exogenous application of cyclo(L-Ala-Gly) to plants (i.e., 21 DAI to BP01R2), the 10 ppm (C10) nor 100 ppm (C100) supplementation contributed to the main root elongation and lateral root development but not to the growth of shoots. Still, apparent contributions on salinity stress alleviation (*i.e.,* vigorous growth of seedlings and the alleviation of leaf chlorosis and adaxial curling) were observed under 50 mM NaCl condition (Fig. 5B).

**Fig. 5.**
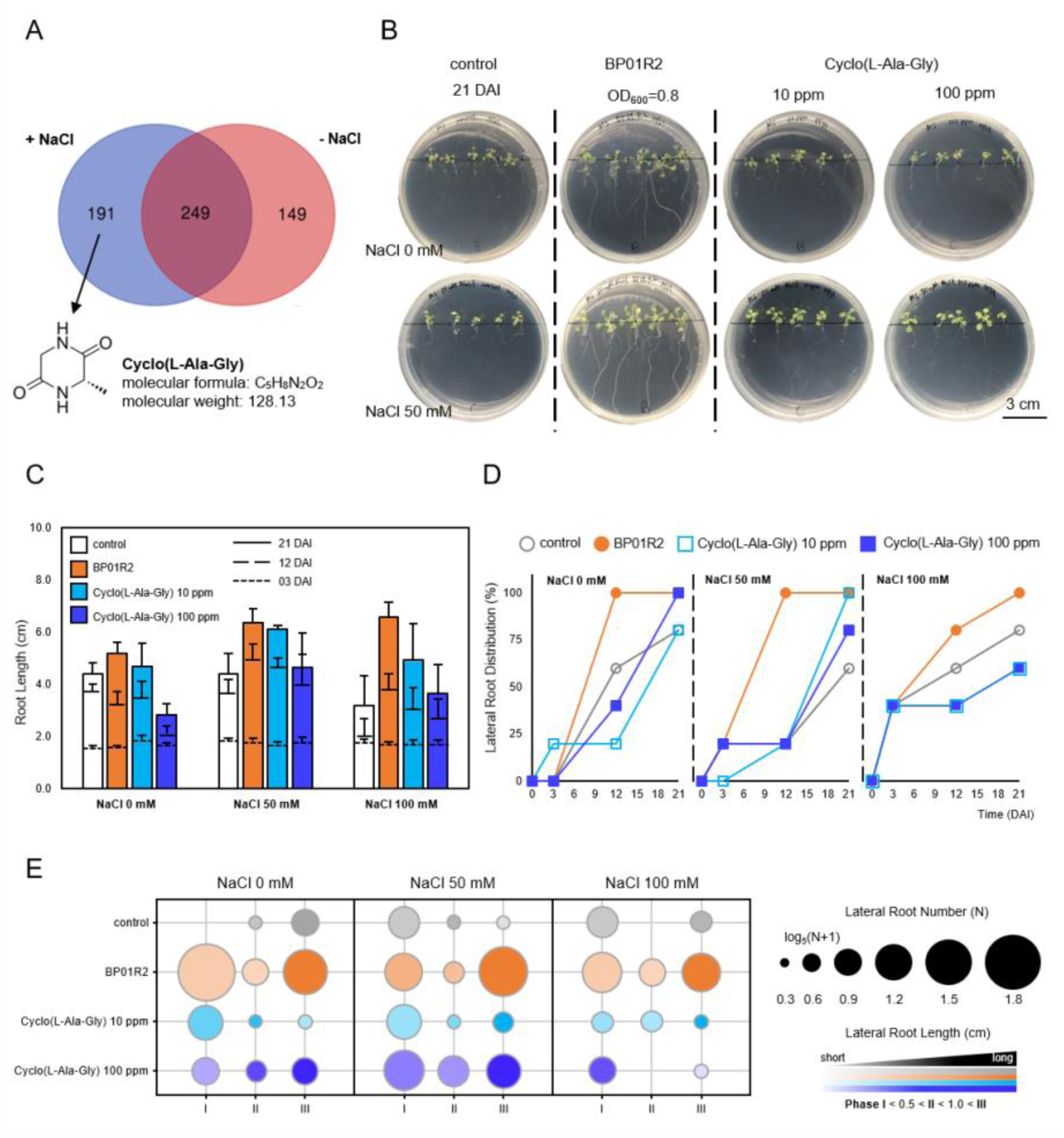
Metabolomic analysis of BP01R2 and the biostimulant candidate mitigating *in planta* salinity stress. **A** Cyclo(L-Ala-Gly) was identified under NaCl-specific bacterial growth conditions. **B** Plant growth differed from that with BP01R2 and cyclo(L-Ala-Gly) under salinity stress. The lower and upper four panels are the plants grown with 50 mM or 0 mM NaCl, respectively. CK: control check; DAI: days after inoculation. **C** main root length. The round dotted line, the dashed line, and the solid line indicate 03, 12 and 21 DAI. **D** Lateral root emergence ratio between 03, 12 and 21 DAI. The open and closed marks of the circle and square indicate the control, BP01R2, cyclo(L-Ala-Gly) 10 ppm and cyclo(L-Ala-Gly) 100 ppm, respectively. **E** The quant and length distribution of lateral roots under different phases. The greater circle size refers to the greater lateral root number. Lateral root developmental phases I, II and III indicate the lateral root lengths <0.5 cm, between 0.5 and 1.0 cm and >1.0 cm and correspond to the colour keys. Data are collected from five independent seedlings with three technical repeats and calculated as mean±SEM.

The decrease in plant main root length was abated by BP01R2, C10 and C100 application under 50 and 100 mM NaCl. For normal plants, much longer main root was observed in BP01R2 and C10 compared to CK. For plants grown under 100 mM NaCl conditions, all treatments showed longer main roots than the CK (Fig. 5C). The lateral root formation ratio of BP01R2 and C100 was 100% compared to 80% of CK and C10 under 0 mM NaCl; the ratio of all treatments was greater than the 60% for CK under 50 mM NaCl; the ratio of C10 and C100 is less than CK, but the BP01R2 remained at 100% under NaCl-free condition (Fig. 5D). The lateral root number of all treatments was more than the CK under 0 mM and 50 mM NaCl, while only the number for BP01R2 was greater than that for CK under 100 mM NaCl. All lateral root were classified into three different developmental phases (phases PI, PII, and PIII indicate the lateral root lengths <0.5 cm, between 0.5 and 1.0 cm and >1.0 cm, respectively). The lateral root length of C100 in PII and PIII was longer than that in CK under 0 mm NaCl; both C10 and C100 could improve longer lateral root development in different phases under 50 mM NaCl; however, both C10 and C100 seem not to contribute much to lateral root development (Fig. 5E). These results suggested that cyclo(L-Ala-Gly), along with other metabolites, may contribute to BP01R2 in alleviating in planta salinity stress and improving plant growth. This indicated its nature as a novel biostimulant, which has already been patented (Taiwan Intellectual Property Office assigned number I684411).

### Genome assembly and comparative genomics of BP01R2 and its relatives

To investigate the key genes for functional characterisation and evolution, we sequenced the genome of strain BP01R2. In total, 2 x 6,486,500 (∼2.0 Gb) of Illumina and 26,722 (∼0.4 Gb; N_50_: 13,469 bp) of trimmed ONT raw reads were obtained for hybrid *de novo* genome assembly. The Illumina and ONT reads provided 349.5X and 66.7X coverage, respectively. The assembly result indicated that BP01R2 has one circular chromosome (5,228,948 bp) with 38.1% GC content and six plasmids with sizes ranging from 5,296 bp to 134,664 bp. For the completeness evaluation, 449 complete BUSCOs (99.8%), 1 fragmented BUSCO (0.2%), and no missing BUSCOs (0.0%) were found within this assembly, which are consistent with the expectation that circular chromosomal contig represents the complete genome assembly. The annotation contains 15 complete sets of rRNA genes and 2 additional 5S rRNA genes, 157 tRNA genes, 8 ncRNAs, 5,551 protein-coding genes, and 91 pseudogenes (Table 1). To investigate the genes associated with MPEI, genes involved in (a) plant growth promotion and rhizosphere competence, (b) salinity stress alleviation and (c) carbon and oxygen limitation adaptation were identified and discussed further for their potential contribution to the symbiotic relationships (Figs 6, 7, S19).

**Fig. 6.**
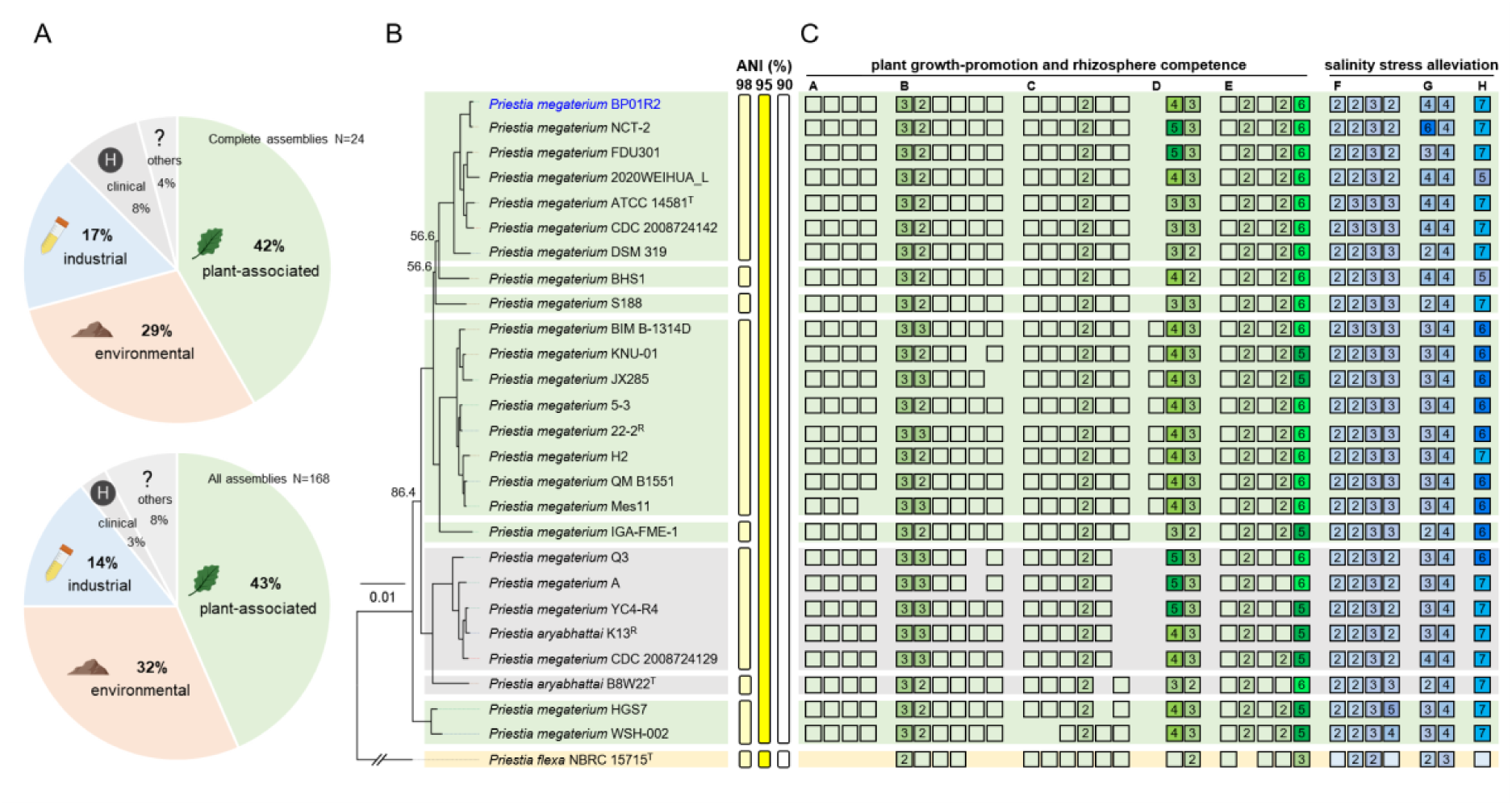
Evolutionary relationships among *Priestia megaterium* BP01R2 and its relatives. **A** Source origin distribution of all available *Priestia megaterium* genomes. **B** Phylogeny of strain BP01R2 and its relatives. The maximum likelihood phylogeny is based on the concatenated alignment of 2,155 single-copy genes shared by all strains analysed (147,671 aligned amino acid sites). The bootstrap values of those three internal nodes with < 99% were labelled. *Priestia flexa* NBRC 15715^T^ is included as the outgroup. The superscripts ‘T’ and ‘R’ following the strain names indicate type strains and strains assigned as NCBI representative genomes, respectively. Strains with complete genomes are presented in bold, and the strain BP01R2 reported in this work is highlighted in blue. Information to the right of the phylogenic tree shows the grouping of genomes according to different cut-off values of genome-wide average nucleotide identity (ANI). **C** Gene content analysis. Empty box and box with numbers inside indicate single-copy and multiple-copy of the targeted genes, respectively. Different genes are clustered in groups: A, nitrogen assimilation and reduction (*nasA-E*); B, phosphate solubilization and mineralization (*gcd, phoP, phoR, phoAB, phoD* and *ppx*); C, L-tryptophane and indole synthesis (*trpA-F*); D, indole-3-acetic acid synthesis and transporter (*gatA* and an AEC family transporter encoded gene); E, acetoin and butanediol synthesis (*alsD, alsS, ilvH* and *ilvG*); F, spermidine synthesis (*speAB* and *speDE*); G, superoxide dismutase and catalase (*sodACF and cat*); H, ferredoxin (*fer*) related. See the gene locus accession details in Table S7. The icons were created with BioRender.com.

**Table 1.** Genomic features of *P. megaterium* strain BP01R2.

| Features | Chromosome | pBP01R2a | pBP01R2b | pBP01R2c | pBP01R2d | pBP01R2e | pBP01R2f |
| --- | --- | --- | --- | --- | --- | --- | --- |
| Accession | CP092387.1 | CP092388.1 | CP092389.1 | CP092390.1 | CP092391.1 | CP092392.1 | CP092393.1 |
| Size (bp) | 5,228,948 | 134,664 | 133,343 | 50,994 | 39,414 | 12,668 | 5,296 |
| GC content (%) | 38 | 34 | 34.5 | 35.5 | 36.5 | 34 | 36 |
| Protein-coding genes | 5231 | 108 | 119 | 44 | 31 | 10 | 8 |
| Pseudogenes | 82 | 3 | 4 | 1 | 0 | 1 | 0 |
| Ribosome RNA genes | 40 | 0 | 0 | 4 | 3 | 0 | 0 |
| Transfer RNA genes | 121 | 0 | 1 | 17 | 18 | 0 | 0 |
| Non-coding RNA genes | 8 | 0 | 0 | 0 | 0 | 0 | 0 |

For the species assignment, we calculated the genome-wide average nucleotide identity (ANI) of BP01R2 and its relatives (Table S1). Additionally, 2,155 single-copy genes shared by all strains were identified and used to produce a concatenated alignment containing 147,671 aligned amino acid sites for maximum likelihood phylogeny construction. The results showed that BP01R2 is closely related to *P. megaterium* NCT-2 and all of the ingroup genomes share > 95% ANI threshold recommended for bacterial species delineation (64) (Fig. 6B). Based on this, strain BP01R2 was identified as *P. megaterium*. We also examined the BioProject and BioSample records of all *P. megaterium* available from NCBI and checked their source origin distribution. As results show, over half of *P. megaterium* were isolated from plant-associated or environmental sources (Fig. 6A). Many of them were reported as beneficial endophytes, PGPB, or originally inhabited in alkaline and hypersaline environments (*e.g.*, salt marshes, alkaline lakes, potash salt dumps, etc.), suggesting their immense potential in dealing with such abiotic stresses (79–81).

For focused gene content investigation, we examined the genes related to MPMI and salinity stress alleviation (Fig. 6C). BP01R2 encodes 180 proteins commonly related to nutrient acquisition, phytohormone production, rhizosphere competence, and ability against abiotic stress in host plants (Fig. S19, Table S7). Genes encoding ABC transporters (Fig. S20) and two-component systems (Fig. S21) are enriched in the BP01R2 genome and likely related to processing environmental information. For rhizosphere competence, five of the *nar*, *nas* and *nir* genes related to nitrogen assimilation and reduction (82,83), as well as a *gcd*, a *ppx* and nine *pho* genes regulating phosphate solubilisation, transport and assimilation (84,85) were found. Additionally, a total of thirty flagellar biosynthesis and assembly associated protein encoding genes (*i.e.*, *flh*, *fli*, *flg* and *mot* genes), eight *che* genes related to bacterial chemotaxis, and three *als*, three *ilv* and six *bdh* genes related to bacterial PGP volatiles (*e.g.*, acetoin and butanediol) (86–88) were present within this genome (Table S7).

For plant-growth improvement, we found eleven genes involved in the indole-3-acetamide (IAM) pathway for indole-3-acetic acid (IAA) biosynthesis from tryptophan (87,89), which was consistent with the metabolomic results for IAA production. Moreover, four auxin efflux carrier (AEC) family transporter protein-encoding genes (90,91), and ten genes encoding siderophore synthesis/Fe-uptake proteins possibly made BP01R2 capable of producing siderophores to assist host plants in chelating ferric ions under iron starvation (83,92) were identified (Fig. 6C, Table S7). Concerning the usual hypoxia and hyper-alkaline/saline condition in a typical niche of BP01R2 and its plant host, abundant genes involved in reductive tricarboxylic acid (rTCA) pathway (72,73), reductive glycine (rGly) pathway (74) and anaerobic respiration metabolism were characterised (Fig. S19, Table S7). Also, four *sod* genes encoding superoxide dismutase family proteins, five *cat* genes encoding catalase proteins, and nine *spe* genes involved in spermidine synthesis related to *in planta* salinity stress alleviation (93) were also found in this genome (Fig. 6C, Table S7).

### Mixotrophic characteristics identification

The BP01R2 genome contains the general nitrite reductase genes *nirBD*, which are required for nitrate ammonification (94), a lactate dehydrogenase gene *ldh*, which primarily mediates the NAD^+^ regeneration during lactate fermentation (95) as well as the required genes involved in the other conversion processes, such as the pyruvate-acetoin-2,3-butanediol and pyruvate-acetyl-CoA-acetate reaction (71,96) (Fig. 7A, S19, Table S7). Meanwhile, the anaerobic regulation related *resA-E* operon encoded (97,98) and *fnr* gene were also found (99) (Table S7). Notably, two nonspontaneous bacterial carbon fixation strategies were present in this genome, *i.e.*, the rGly pathway (74) and the rTCA cycle (72,73) (Figs 7A, S19, Table S7). To estimate the anaerobic and autotrophic availability of BP01R2, we incubated it under oxygen and carbon resource starvation conditions by mimicking the original habitat of its photoautotrophic host. The results showed that BP01R2 grow under anaerobic conditions (90% H_2_ plus 10% CO_2_) (Fig. 7B), and a high correlation (R^2^=0.95) were found between the extra glucose supplementation and the maximum bacterial growing capability (Fig. 7C), which indicates the availability of BP01R2 in acquiring organic carbon resources for growth under an anaerobic condition.

**Fig. 7.**
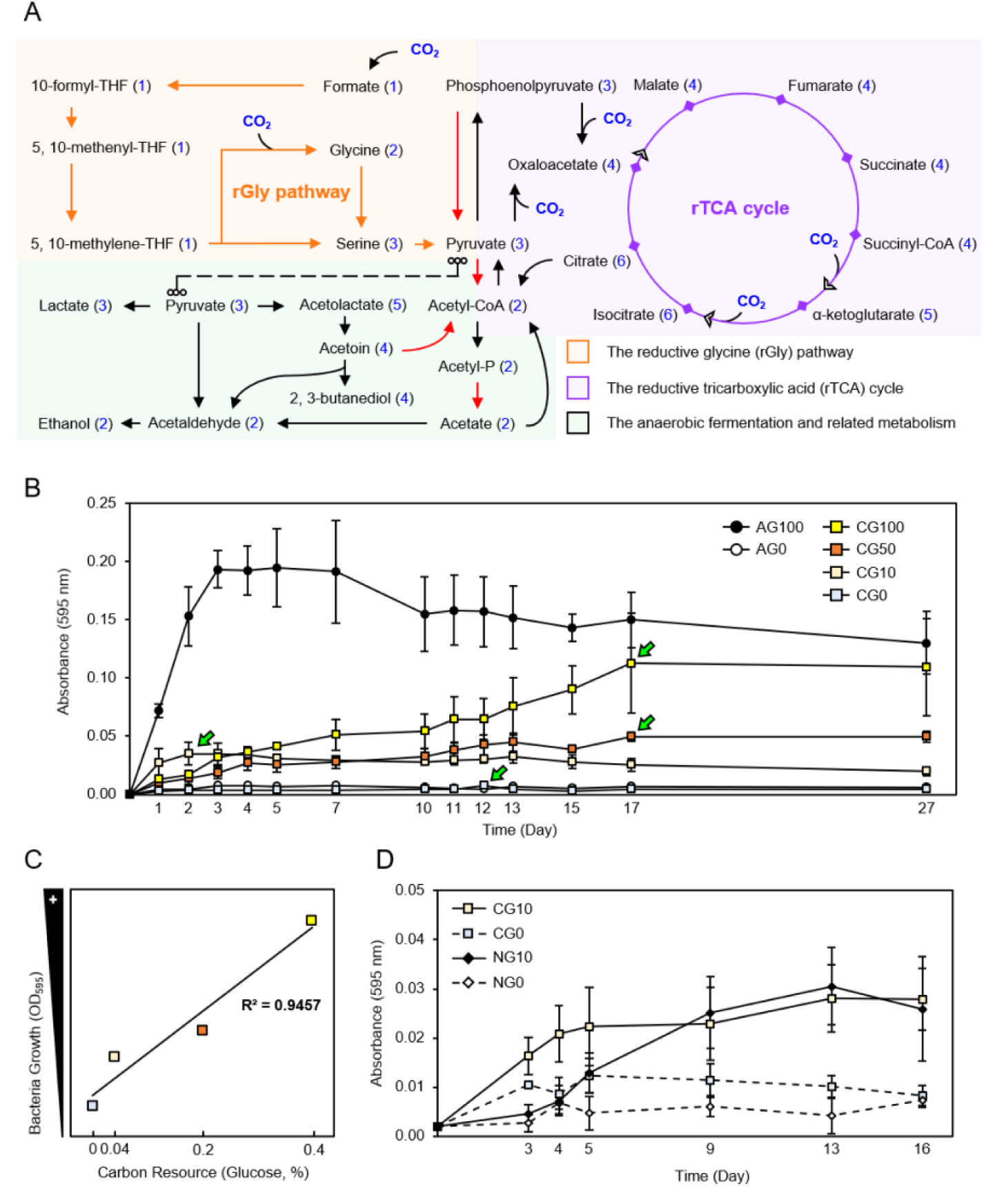
Tropism analyses on BP01R2. **A** The simplified autotrophic (orange and purple panels) and anaerobic (green panel) related metabolism identified within the BP01R2 genome. All of the gene names and locus accession details are listed in Table S7. Carbon dioxide (CO_2_) influx and the number of carbon atoms in each molecule are highlighted in blue; the reactions involved in energy generation are marked as red arrows. **B D** Bacterial growth curves under different carbon resources (G0, G10, G50 and G100 indicate 0%, 0.04%, 0.2% and 0.4% glucose, respectively) and tube headspace gas compositions (A, C and N indicate sterilised air, 90% H_2_ plus 10% CO_2_, and 90% H_2_ plus 10% N_2_, respectively). **C** Coefficient relationship between carbon resources and bacterial growth under anaerobic conditions. X-axis, glucose content; Y-axis, the maxima (indicated by the fluorescent green arrows) of the bacterial growth curves in **B**. All data points are collected from three independent biological repeats and shown as mean±SEM.

Furthermore, we were interested in whether BP01R2 use CO_2_ as a sole carbon resource to grow under anaerobic conditions, as a chemolithoautotroph. The testing glucose supplantation was 0.04% and 0%, and N_2_ was used to replace the CO_2_ in anaerobic gas. The results confirmed that BP01R2 can use CO_2_ for growth either with or without extra 0.04% glucose (Fig. 7D). Interestingly, bacteria grew significantly better with CO_2_ than N_2_ on the 3rd day (CG0 vs NG0, p=0.05; CG10 vs NG10, p=0.03), and this advantage was kept when growing without glucose till the 16th day. A similar pattern was found in the 0.04% glucose conditions before the 9th day, and a higher growth rate was observed in those CO_2_ present treatments; however, no significance was found in bacterial maximum growth capacity compared to the N_2_ conditions after then. Accordingly, BP01R2 was identified as a facultative aerobic mixotroph that can use organic (glucose) or inorganic (CO_2_) carbon resources to grow.

## Discussion

### BP01R2 and cyclo(L-Ala-Gly) as versatile endophytic biostimulants

Auxin is well known as a crucial instructor to root/shoot morphogenesis and development by converging multiple phytohormones and signalling pathways in plants (100–102). In this work, several Indole-3-acetic acid (IAA)-producing-related and auxin efflux carrier (AEC) family transporter genes (Table S7) and the IAA production (Fig. S18, Tables S3, S4) of BP01R2 were confirmed. Besides, some plants’ growth-promoting bacterial volatiles and (lateral) root-inducing biostimulants (Table S4, S5) and over 40 genes related to bacterial flagella and chemotaxis (Table S7) were identified in the BP01R2 genome. Some endophytes have bacterial flagella and produce biostimulants, which cause different repulsive or attractive compounds and quorum sensing. The contact between endophytes and plant exudates occurs accordingly, and chemotaxis driven by flagella also plays an important role in colonisation (92,103–105).

The root structures will change for morphological plasticity preventing salt accumulation from the saline environment while plants suffer the salinity stress (106,107). Besides phytohormones, multiple pathways like glutathione metabolism and ribosome biosynthesis will respond in an opposite pattern than those without stress (108–110). Similar phenomena were also found within BP01R2 inoculated plants (Figs 3, S10-S17), suggesting its benefits to *in planta* salinity stress alleviation. While looking into bacterial metabolomics under salinity stress, we noticed some signals of cyclopeptides; then, we further confirmed one cyclic dipeptide, cyclo(L-Ala-Gly), functions like BP01R2 in plants (Fig. 5). Distinct from our previous work describing the PGP traits and application (26), here we decipher the BP01R2’s genome and metabolome and examine the plant transcriptome to reveal the molecular mechanisms of its MPMI and stress alleviation in the plant. All evidence collectively suggests that BP01R2 improves growth and alleviates salinity stress with beneficial biostimulants and transcriptomic regulation in the host plant (Fig. 8A).

**Fig. 8.**
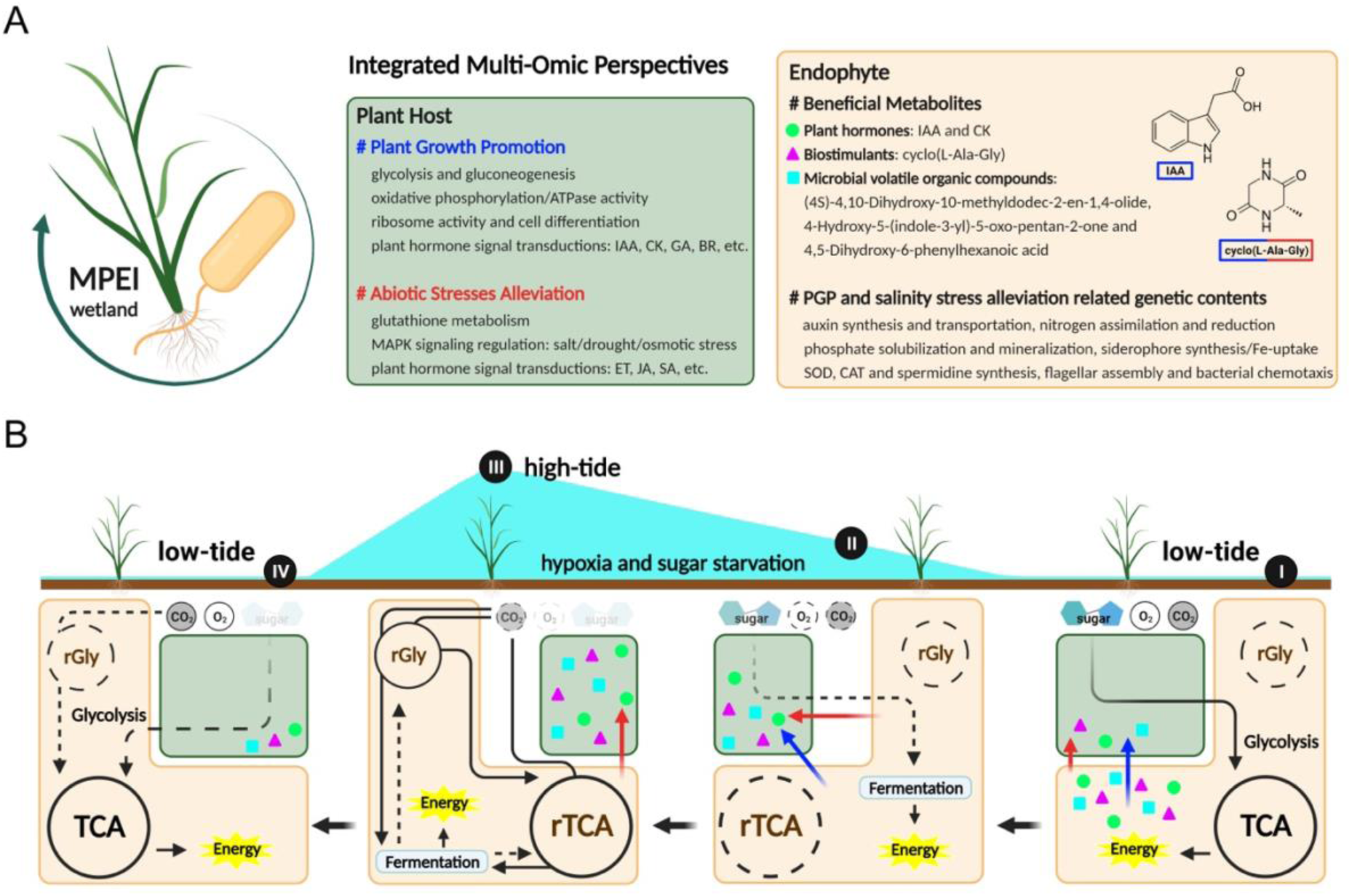
Schematic illustration of molecular plant-endophyte interactions (MPEI) among wetland biomes. **A** Integrated multi-omics perspectives on the symbiotic relationships of wetland plants and endophytes. Green cell indicated those enriched phenotypic and transcriptomic regulations within the plant contributed by the endophyte; orange cell was generated based on metabolomic and genomic data analyses in this work. **B** The dynamic carbon resource acquisition switching of the endophyte may be a niche for host-symbiosis under hypoxia and sugar starvation. For simplicity, the black arrow represents any carbon flux from organic (sugar) or inorganic (CO_2_) carbon sources. Both dashed lines and transparent marks indicate a decreasing level of suitability or availability. Phases I, II, III and IV indicate four timepoints among the tidal dynamics of the wetland biome, and the diagrams below indicate the timepoint-specific MPEI accordingly. Blue and red marks highlight the plant growth promotion and abiotic stress alleviation effects on the plant host, respectively. Brightly coloured symbols indicate the corresponding beneficial metabolites mentioned in the orange cell of **A**. Icons were created with BioRender.com.

Although the potential of CDPs as stress-mitigating biostimulants, especially cyclo(L-Ala-Gly), has been revealed in this work, further investigations are required in future work to decipher its molecular roles in plants. The mechanisms of these compounds in plants remain unclear; one of the reasons is that developing an efficient and specific method for detecting their *in planta* amount and distribution remains a technical difficulty. In summary, this work facilitates our understanding of CDPs and highlights the importance of these potent small compounds that deserve more attention.

### Taxonomy of *Priestia* species

Based on information available from the strains with NCBI BioProject/BioSample information, > 70% of *P. megaterium* strains are associated with plants or are environmental microbes (Fig. 6A) and all have abundant PGPR and stress abatement related genetic components (Fig. 6C), suggesting the potential of this species for agricultural applications. Based on genome-wide ANI analysis and molecular phylogeny of conserved single-copy genes, some of the characterised strains may be mis-classified. Notably, our results showed that the type strain of *P. aryabhattai* shares > 95% ANI with *P. megaterium* (Fig. 6B). To clarify the species assignments, we backtracked the supporting information described for these species delineation (111). However, two conserved signature indels (CSIs) reported by Gupta *et al.* (111) were absent in neither the type strain of *P. megaterium* nor *P. aryabhattai* (Fig. S22), suggesting that these markers are not reliable. Based on the International Code of Nomenclature of Prokaryotes (Principle 8, ‘Each order or taxon of a lower rank with a given circumscription, position, and rank can bear only one correct name, *i.e.*, the earliest that is in accordance with the Rules of this Code’) (112), we recommend to reject the name *Priestia aryabhattai* (Basonym: Bacillus aryabhattai (113)) and temporarily replaced it with *Priestia megaterium* (Basonym: *Bacillus megaterium* (114)). This taxonomic reclassification coincides with the synonymy suggestion by Narsing Rao *et al.* (115). Meanwhile, further confirmation of the correctness and suitability of the name *Priestia* gen. nov. is required to avoid incorrect uses.

### Insights into symbiosis among wetland MPEI

As mentioned, BP01R2 originally inhabited salt marsh plants that grow with continuously high salinity and periodic hypoxia (Fig. 1A). When plants suffer abiotic stress, *e.g.*, salinity stress, drought stress and osmotic stress, the transmissible sugar in plants tends to accumulate in the vacuole for water pressure manipulation or recycled into storage forms of sugar (*e.g.*, starch, cellulose, lignin, etc.), and microorganisms living in plant endosphere may suffer from *in planta* carbon resource starvation (116–118). There is consensus that efficient carbon resource transmissions (*i.e.*, sugars, amino acids, other organic matters, etc.) from plant phloem and root exudates to symbionts are important (119,120). Therefore, addressing this routine multi-stress condition is important for symbiosis maintenance among endophytes. The genomics (Figs 7A, S19, Table S7) and tropism analysis on BP01R2 reveal its anaerobic and autotrophic metabolisms availability and mixotrophic traits, allowing it to grow under the mimic condition of wetland plant suffering multi-stresses of sugar starvation and hypoxia *in vitro* (Figs 7B-D).

Collectively, in this study, we infer that the anaerobic and autotrophic characteristics of the endophyte may reduce the carbon source demand from the host under hypoxia stress, such as during the high-tide period in wetlands (from Phase I to Phase III, Fig. 8B), *i.e.*, the pathway by which strains gain energy might be able to be switched from consuming sugar and the TCA cycle into anaerobic carbon fixation, such as the rTCA cycle, and fermentation (Phase III, Fig. 8B). These may be a strategy and a niche for symbionts increasing competitiveness for host symbiosis under multi-stress or extreme biome. Herein, based on the multi-omics investigations and dynamic switching of carbon resources and energy acquisition of *P. megaterium* BP01R2, we provide a novel perspective on molecular plant-endophyte interactions (MPEI) in the wetland biome (Fig. 8). However, the coordination of carbon resource acquisition (organic and inorganic) and trophic patterns (heterotrophic and autotrophic) among MPMI remain obscure. Multi-dimensional gene regulation research on plant-microbe interactions (121–123) and tracing fluxes of carbon resources among bacteria and plant hosts (124) are necessary for future work.

## Conclusions

Beyond uncovering the versatile potential of *P. megaterium* BP01R2 and cyclo(L-Ala-Gly) for future agriculture, our multi-omics and bacterial tropism results provide new insights into the unclear mechanism of trophic homeostasis among MPEI. Multitudes of microorganisms colonise diverse compartments of healthy plants, and their importance in the modulation of commensal or mutual relationships has been widely discussed (120,125,126). How bacterial tropism shapes the endophytes’ autotrophic and energy metabolisms, either in transcriptomic or metabolic dynamics, to maintain their competitiveness among host-microbe symbiosis systems remains an outstanding issue. The symbiosis constructed of mixoautotroph and photoautotroph reported here provides a preliminary but novel perspective and untapped opportunities to understand the enigmatic plant-endophyte symbiotic relationships in MPMI.

## Data Availability Statement

The complete genome sequence of *Priestia megaterium* BP01R2 has been deposited in GenBank under the accession numbers CP092387 (chromosome) and CP092388-93 (plasmids). The genome and RNA sequencing project and the associated raw reads were deposited in the NCBI under BioProject PRJNA806882 and PRJNA818431, respectively.

## Acknowledgements

The Illumina library preparation and sequencing services of the bacterial genome were provided by the Genomics BioSci & Tech Co., Ltd. (Taipei, Taiwan) and Genomic Technology Core (Institute of Plant and Microbial Biology, Academia Sinica); the ONT sequencing services were provided by the Inong Agriculture Co., Ltd. (Taipei, Taiwan). The RNAseq services were provided by Welgene Biotech Co., Ltd. (Taipei, Taiwan). We also thank the support from the Taichung District Agricultural Research and Extension Station, Council of Agriculture, Executive Yuan of Taiwan.

## Funding

The funding for this project was provided by the National Science and Technology Council of Taiwan (MOST 109-2321-B-005-025; MOST 110-2321-B-005-008), the Ministry of Agriculture of Taiwan (110AS-1.6.1-BQ-B3; 111AS-1.6.1-ST-a7) and the Ministry of Education of Taiwan (The Higher Education Sprout Project) to Chieh-Chen Huang, and Academia Sinica to Chih-Horng Kuo. The funders of this work had no role in study design, data collection and interpretation, or the decision for publication.

## Competing Interests

The concepts of the paper are shared with a patent filed by the National Chung Hsing University: Taiwan Patent No. I684411, on which Pin-Hsien Yeh, Ying-Ning Ho and Chieh-Chen Huang are named inventors.

## References

1. Policy Institute IF. 2022 Global food policy report: climate change and food systems [Internet]. Washington, DC: International Food Policy Research Institute; 2022 [cited 2022 Jul 9]. Available from: https://ebrary.ifpri.org/digital/collection/p15738coll2/id/135889

2. Stads GJ, Wiebe KD, Nin-Pratt A, Sulser TB, Benfica R, Reda F, et al. Research for the future: investments for efficiency, sustainability, and equity [Internet]. Washington, DC: International Food Policy Research Institute; 2022 [cited 2022 Jul 9]. Available from: https://ebrary.ifpri.org/digital/collection/p15738coll2/id/135885

3. Swinnen J, Arndt C, Vos R. Climate change and food systems: transforming food systems for adaptation, mitigation, and resilience [Internet]. Washington, DC: International Food Policy Research Institute; 2022 [cited 2022 Jul 9]. Available from: https://ebrary.ifpri.org/digital/collection/p15738coll2/id/135884

4. Ha-Tran DM, Nguyen TTM, Hung SH, Huang E, Huang CC. Roles of plant growth-promoting rhizobacteria (PGPR) in stimulating salinity stress defense in plants: a review. International Journal of Molecular Sciences. 2021 Jan;22(6):3154.

5. Hung SHW, Huang TC, Lai YC, Wu IC, Liu CH, Huarng YF, et al. Endophytic biostimulants for smart agriculture: Burkholderia seminalis 869T2 benefits heading leafy vegetables in-field management in Taiwan. Agronomy. 2023 Apr;13(4):967.

6. Rosenblueth M, Martínez-Romero E. Bacterial endophytes and their interactions with hosts. Mol Plant Microbe Interact. 2006 Aug;19(8):827–37.

7. White JF, Kingsley KL, Zhang Q, Verma R, Obi N, Dvinskikh S, et al. Review: endophytic microbes and their potential applications in crop management. Pest Manag Sci. 2019 Oct;75(10):2558–65.

8. Elsheikh EAE, El-Keblawy A, Mosa KA, Okoh AI, Saadoun I. Role of Endophytes and Rhizosphere Microbes in Promoting the Invasion of Exotic Plants in Arid and Semi-Arid Areas: A Review. Sustainability. 2021 Jan;13(23):13081.

9. Solanki MK. Phytobiomes: Current Insights And Future Vistas [Internet]. Kashyap PL, Kumari B, editors. Toronto, ON: Springer Nature; 2021 [cited 2022 Jul 9]. Available from: https://www.chapters.indigo.ca/en-ca/books/phytobiomes-current-insights-and-future/9789811531538-item.html

10. Johnston-Monje D, Raizada MN. Conservation and diversity of seed associated endophytes in Zea across boundaries of evolution, ethnography and ecology. PLoS One. 2011;6(6):e20396.

11. Cipriano MAP, Freitas-Iório R de P, Dimitrov MR, de Andrade SAL, Kuramae EE, Silveira APD da. Plant-Growth Endophytic Bacteria Improve Nutrient Use Efficiency and Modulate Foliar N-Metabolites in Sugarcane Seedling. Microorganisms. 2021 Feb 25;9(3):479.

12. García-Latorre C, Rodrigo S, Santamaria O. Effect of fungal endophytes on plant growth and nutrient uptake in Trifolium subterraneum and Poa pratensis as affected by plant host specificity. Mycol Progress. 2021 Sep 1;20(9):1217– 31.

13. Mishra S, Priyanka, Sharma S. Metabolomic Insights Into Endophyte-Derived Bioactive Compounds. Frontiers in Microbiology [Internet]. 2022 [cited 2022 Jul 9];13. Available from: https://www.frontiersin.org/articles/10.3389/fmicb.2022.835931

14. Verma SK, Sahu PK, Kumar K, Pal G, Gond SK, Kharwar RN, et al. Endophyte roles in nutrient acquisition, root system architecture development and oxidative stress tolerance. J Appl Microbiol. 2021 Nov;131(5):2161–77.

15. Verma H, Kumar D, Kumar V, Kumari M, Singh SK, Sharma VK, et al. The Potential Application of Endophytes in Management of Stress from Drought and Salinity in Crop Plants. Microorganisms. 2021 Aug 13;9(8):1729.

16. Davy AJ, Brown MJH, Mossman HL, Grant A. Colonization of a newly developing salt marsh: disentangling independent effects of elevation and redox potential on halophytes: Elevation and redox potential effects on halophytes. Journal of Ecology. 2011 Nov;99(6):1350–7.

17. Janousek CN, Folger CL. Variation in tidal wetland plant diversity and composition within and among coastal estuaries: assessing the relative importance of environmental gradients. Halvorsen R, editor. J Veg Sci. 2014 Mar;25(2):534–45.

18. Veldhuis ER, Schrama M, Staal M, Elzenga JTM. Plant Stress-Tolerance Traits Predict Salt Marsh Vegetation Patterning. Frontiers in Marine Science. 2019;5:501.

19. Syranidou E, Christofilopoulos S, Gkavrou G, Thijs S, Weyens N, Vangronsveld J, et al. Exploitation of Endophytic Bacteria to Enhance the Phytoremediation Potential of the Wetland Helophyte Juncus acutus. Frontiers in Microbiology. 2016;7:1016.

20. Chandra R, Singh K. Phytoremediation of environmental pollutants. In: Chandra R, Dubey NK, Kumar V, editors. Phytoremediation of Environmental Pollutants [Internet]. 1st ed. NY, USA: Taylor & Francis, CRC Press; 2017. p. 42. Available from: https://www.taylorfrancis.com/chapters/edit/10.1201/9781315161549-12/endophytic-bacterial-diversity-roots-wetland-plants-potential-enhancing-phytoremediation-environmental-pollutants-ram-chandra-kshitij-singh

21. Ashraf S, Afzal M, Rehman K, Tahseen R, Naveed M, Zahir ZA. Enhanced remediation of tannery effluent in constructed wetlands augmented with endophytic bacteria. dwt. 2018;102:93–100.

22. Singh T, Awasthi G, Tiwari Y. Recruiting endophytic bacteria of wetland plants to phytoremediate organic pollutants. Int J Environ Sci Technol [Internet]. 2021 Jun 16 [cited 2022 Jul 9]; Available from: https://link.springer.com/10.1007/s13762-021-03476-y

23. Ho YN, Chiang HM, Chao CP, Su CC, Hsu HF, Guo CT, et al. In planta biocontrol of soilborne Fusarium wilt of banana through a plant endophytic bacterium, Burkholderia cenocepacia 869T2. Plant Soil. 2015 Feb 1;387(1):295–306.

24. Nguyen BAT, Hsieh JL, Lo SC, Wang SY, Hung CH, Huang E, et al. Biodegradation of dioxins by Burkholderia cenocepacia strain 869T2: role of 2-haloacid dehalogenase. Journal of Hazardous Materials. 2021 Jan 5;401:123347.

25. Hwang HH, Chien PR, Huang FC, Hung SH, Kuo CH, Deng WL, et al. A plant endophytic bacterium, Burkholderia seminalis strain 869T2, promotes plant growth in Arabidopsis, Pak Choi, Chinese amaranth, lettuces, and other vegetables. Microorganisms. 2021 Aug 10;9(8):1703.

26. Hwang HH, Chien PR, Huang FC, Yeh PH, Hung SHW, Deng WL, et al. A plant endophytic bacterium Priestia megaterium strainBP-R2 isolated from the halophyte Bolboschoenus planiculmis enhances plant growth under salt and drought stresses. Microorganisms. 2022 Oct;10(10):2047.

27. Otieno N, Lally R, Kiwanuka S, Lloyd A, Ryan D, Germaine K, et al. Plant growth promotion induced by phosphate solubilizing endophytic Pseudomonas isolates. Frontiers in Microbiology. 2015;6:745.

28. Wang X, Li Y, Zhang X, Lai D, Zhou L. Structural Diversity and Biological Activities of the Cyclodipeptides from Fungi. Molecules. 2017 Dec;22(12):2026.

29. Hung SHW, Chiu MC, Huang CC, Kuo CH. Complete genome sequence of Curtobacterium sp. C1, a beneficial endophyte with the potential for in-plant salinity stress alleviation. MPMI. 2022 Jul 12;35(8):731–5.

30. Lo SC, Tsai SY, Chang WH, Wu IC, Sou NL, Hung SHW, et al. Characterization of the Pyrroloquinoline Quinone Producing Rhodopseudomonas palustris as a Plant Growth-Promoting Bacterium under Photoautotrophic and Photoheterotrophic Culture Conditions. International Journal of Molecular Sciences. 2023 Jan;24(18):14080.

31. Degrassi G, Aguilar C, Bosco M, Zahariev S, Pongor S, Venturi V. Plant Growth-Promoting Pseudomonas putida WCS358 Produces and Secretes Four Cyclic Dipeptides: Cross-Talk with Quorum Sensing Bacterial Sensors. Curr Microbiol. 2002 Oct 1;45(4):250–4.

32. Kumar NS, Mohandas C. Antimycobacterial activity of cyclic dipeptides isolated from Bacillus sp. N strain associated with entomopathogenic nematode. Pharmaceutical Biology. 2014 Jan 1;52(1):91–6.

33. Ortiz A, Sansinenea E. Cyclic Dipeptides: Secondary Metabolites Isolated from Different Microorganisms with Diverse Biological Activities. Curr Med Chem. 2017;24(25):2773–80.

34. Prasad C. Bioactive cyclic dipeptides. Peptides. 1995 Jan;16(1):151–64.

35. Wattana-Amorn P, Charoenwongsa W, Williams C, Crump MP, Apichaisataienchote B. Antibacterial activity of cyclo(L -Pro- L -Tyr) and cyclo(D -Pro- L -Tyr) from *Streptomyces* sp. strain 22-4 against phytopathogenic bacteria. Natural Product Research. 2016 Sep 1;30(17):1980– 3.

36. Kurohashi M, Nakamura K, Ienaga K. Agricultural and horticultural compositions inducing resistance in plants against salt- and water-stresses [Internet]. Hyogo, Japan; EP0331641A2, 1989 [cited 2022 Jul 10]. Available from: https://patents.google.com/patent/EP0331641A2/en

37. Edgar RC. MUSCLE: multiple sequence alignment with high accuracy and high throughput. Nucleic Acids Research. 2004 Mar 8;32(5):1792–7.

38. Guindon S, Gascuel O. A simple, fast, and accurate algorithm to estimate large phylogenies by maximum likelihood. Rannala B, editor. Systematic Biology. 2003 Oct 1;52(5):696–704.

39. Felsenstein J. PHYLIP - phylogeny inference package (version 3.2). Cladistics. 1989;5:164–6.

40. Rivero L, Scholl R, Holomuzki N, Crist D, Grotewold E, Brkljacic J. Handling Arabidopsis plants: growth, preservation of seeds, transformation, and genetic crosses. In: Sanchez-Serrano JJ, Salinas J, editors. Arabidopsis protocols [Internet]. Totowa, NJ: Humana Press; 2014 [cited 2022 Jul 10]. p. 3–25. Available from: http://link.springer.com/10.1007/978-1-62703-580-4_1

41. Tsai YH. Transcriptomic analysis on adaptive immunity of Pei-Chiao (Musa spp., Cavendish AAA) against Fusarium wilt caused by Fusarium oxysporum f. sp. cubense tropical race4. [MS thesis]. [Taichung, Taiwan]: National Chung Hsing University; 2019.

42. Yu MY. A study of pyrroloquinoline quinone on Pei-Chiao (Musa spp.) growth and biocontrol improvement [MS thesis]. [Taichung, Taiwan]: National Chung Hsing University; 2019.

43. Berardini TZ, Reiser L, Li D, Mezheritsky Y, Muller R, Strait E, et al. The arabidopsis information resource: Making and mining the “gold standard” annotated reference plant genome: Tair: Making and Mining the “Gold Standard” Plant Genome. genesis. 2015 Aug;53(8):474–85.

44. Wagner GP, Kin K, Lynch VJ. Measurement of mRNA abundance using RNA-seq data: RPKM measure is inconsistent among samples. Theory Biosci. 2012 Dec;131(4):281–5.

45. Love MI, Huber W, Anders S. Moderated estimation of fold change and dispersion for RNA-seq data with DESeq2. Genome Biology. 2014 Dec 5;15(12):550.

46. Pertea M, Kim D, Pertea GM, Leek JT, Salzberg SL. Transcript-level expression analysis of RNA-seq experiments with HISAT, StringTie and Ballgown. Nat Protoc. 2016 Sep;11(9):1650–67.

47. Subramanian A, Tamayo P, Mootha VK, Mukherjee S, Ebert BL, Gillette MA, et al. Gene set enrichment analysis: A knowledge-based approach for interpreting genome-wide expression profiles. Proc Natl Acad Sci USA. 2005 Oct 25;102(43):15545–50.

48. Mi H, Muruganujan A, Ebert D, Huang X, Thomas PD. PANTHER version 14: more genomes, a new PANTHER GO-slim and improvements in enrichment analysis tools. Nucleic Acids Research. 2019 Jan 8;47(D1):D419–26.

49. Kanehisa M, Goto S. KEGG: Kyoto encyclopedia of genes and genomes. Nucleic Acids Res. 2000 Jan 1;28(1):27–30.

50. Chambers MC, Maclean B, Burke R, Amodei D, Ruderman DL, Neumann S, et al. A cross-platform toolkit for mass spectrometry and proteomics. Nat Biotechnol. 2012 Oct;30(10):918–20.

51. Pluskal T, Castillo S, Villar-Briones A, Orešič M. MZmine 2: modular framework for processing, visualizing, and analyzing mass spectrometry-based molecular profile data. BMC Bioinformatics. 2010 Dec;11(1):395.

52. Verkh Y, Rozman M, Petrovic M. Extraction and cleansing of data for a non-targeted analysis of high-resolution mass spectrometry data of wastewater. MethodsX. 2018;5:395–402.

53. Blunt JW, Munro MHG, Laatsch H. AntiMarin database. University of Canterbury, Christchurch, New Zealand; University of Gottingen, Gottingen, Germany [Internet]. 2006; Available from: https://scholar.google.com/scholar_lookup?title=AntiMarin+Database&publication_year=2006&

54. Hung SHW, Wu IC, Huang CC, Kuo CH. Complete genome sequence of Erythrobacteraceae bacterium WH01K, a strain isolated from corals (Acropora sp.) in Taiwan. Microbiology Resource Announcements. 2023 Nov 7;0(0):e00830–23.

55. Wick RR, Judd LM, Gorrie CL, Holt KE. Unicycler: resolving bacterial genome assemblies from short and long sequencing reads. PLoS Comput Biol. 2017 Jun 8;13(6):e1005595.

56. Li H, Durbin R. Fast and accurate short read alignment with Burrows-Wheeler transform. Bioinformatics. 2009 Jul 15;25(14):1754–60.

57. Li H. Minimap2: pairwise alignment for nucleotide sequences. Birol I, editor. Bioinformatics. 2018 Sep 15;34(18):3094–100.

58. Li H, Handsaker B, Wysoker A, Fennell T, Ruan J, Homer N, et al. The sequence alignment/map format and SAMtools. Bioinformatics. 2009 Aug 15;25(16):2078–9.

59. Robinson JT, Thorvaldsdóttir H, Winckler W, Guttman M, Lander ES, Getz G, et al. Integrative genomics viewer. Nat Biotechnol. 2011 Jan;29(1):24–6.

60. Nishimura O, Hara Y, Kuraku S. Evaluating genome assemblies and gene models using gVolante. Methods Mol Biol. 2019 Jan 1;1962:247–56.

61. Manni M, Berkeley MR, Seppey M, Simão FA, Zdobnov EM. BUSCO update: novel and streamlined workflows along with broader and deeper phylogenetic coverage for scoring of eukaryotic, prokaryotic, and viral genomes. Mol Biol Evol. 2021 Sep 1;38(10):4647–54.

62. Tatusova T, DiCuccio M, Badretdin A, Chetvernin V, Nawrocki EP, Zaslavsky L, et al. NCBI prokaryotic genome annotation pipeline. Nucleic Acids Res. 2016 Aug 19;44(14):6614–24.

63. Aramaki T, Blanc-Mathieu R, Endo H, Ohkubo K, Kanehisa M, Goto S, et al. KofamKOALA: KEGG Ortholog assignment based on profile HMM and adaptive score threshold. Bioinformatics. 2020 Apr 1;36(7):2251–2.

64. Jain C, Rodriguez-R LM, Phillippy AM, Konstantinidis KT, Aluru S. High throughput ANI analysis of 90K prokaryotic genomes reveals clear species boundaries. Nat Commun. 2018 Dec;9(1):5114.

65. Li L, Stoeckert CJ, Roos DS. OrthoMCL: identification of ortholog groups for eukaryotic genomes. Genome Res. 2003 Sep;13(9):2178–89.

66. Dahmani MA, Desrut A, Moumen B, Verdon J, Mermouri L, Kacem M, et al. Unearthing the Plant Growth-Promoting Traits of Bacillus megaterium RmBm31, an Endophytic Bacterium Isolated From Root Nodules of Retama monosperma. Frontiers in Plant Science [Internet]. 2020 [cited 2022 Jul 10];11. Available from: https://www.frontiersin.org/articles/10.3389/fpls.2020.00124

67. Santoyo G, Moreno-Hagelsieb G, del Carmen Orozco-Mosqueda Ma, Glick BR. Plant growth-promoting bacterial endophytes. Microbiological Research. 2016 Feb;183:92–9.

68. Du CX, Fan HF, Guo SR, Tezuka T. Applying Spermidine for Differential Responses of Antioxidant Enzymes in Cucumber Subjected to Short-term Salinity. J Amer Soc Hort Sci. 2010 Jan;135(1):18–24.

69. Li S, Fan C, Li Y, Zhang J, Sun J, Chen Y, et al. Effects of drought and salt-stresses on gene expression in Caragana korshinskii seedlings revealed by RNA-seq. BMC Genomics. 2016 Mar 8;17:200.

70. Saibi W, Brini F. Superoxide dismutase (SOD) and abiotic stress tolerance in plants: An overview. In: Magliozzi S, editor. Superoxide Dismutase: Structure, Synthesis and Applications [Internet]. Hauppauge, NY, USA: Nova Science; 2018. p. 101–42. Available from: https://www.researchgate.net/profile/Faical-Brini/publication/322551748_Superoxide_dismutase_SOD_and_abiotic_stress_tolerance_in_plants_An_overview/links/5a605d21a6fdcc21f487c424/Superoxide-dismutase-SOD-and-abiotic-stress-tolerance-in-plants-An-overview.pdf

71. Cruz Ramos H, Hoffmann T, Marino M, Nedjari H, Presecan-Siedel E, Dreesen O, et al. Fermentative Metabolism of *Bacillus subtilis* : Physiology and Regulation of Gene Expression. J Bacteriol. 2000 Jun;182(11):3072–80.

72. Mall A, Sobotta J, Huber C, Tschirner C, Kowarschik S, Bačnik K, et al. Reversibility of citrate synthase allows autotrophic growth of a thermophilic bacterium. Science. 2018 Feb 2;359(6375):563–7.

73. Nunoura T, Chikaraishi Y, Izaki R, Suwa T, Sato T, Harada T, et al. A primordial and reversible TCA cycle in a facultatively chemolithoautotrophic thermophile. Science. 2018 Feb 2;359(6375):559–63.

74. Sánchez-Andrea I, Guedes IA, Hornung B, Boeren S, Lawson CE, Sousa DZ, et al. The reductive glycine pathway allows autotrophic growth of Desulfovibrio desulfuricans. Nat Commun. 2020 Oct 1;11(1):5090.

75. Cultivation of Anaerobes [Internet]. [cited 2022 May 30]. Available from: https://www.dsmz.de/fileadmin/Bereiche/Microbiology/Dateien/Kultivierungshinweise/englAnaerob.pdf

76. Charles Zaiontz. Real Statistics Resource Pack software [Internet]. 2020. Available from: www.real-statistics.com

77. Lally RD, Galbally P, Moreira AS, Spink J, Ryan D, Germaine KJ, et al. Application of Endophytic Pseudomonas fluorescens and a Bacterial Consortium to Brassica napus Can Increase Plant Height and Biomass under Greenhouse and Field Conditions. Frontiers in Plant Science. 2017;8:2193.

78. Tamošiūnė I, Stanienė G, Haimi P, Stanys V, Rugienius R, Baniulis D. Endophytic Bacillus and Pseudomonas spp. Modulate Apple Shoot Growth, Cellular Redox Balance, and Protein Expression Under in Vitro Conditions. Frontiers in Plant Science. 2018;9:889.

79. Vílchez JI, Tang Q, Kaushal R, Wang W, Lv S, He D, et al. Complete Genome Sequence of Bacillus megaterium Strain TG1-E1, a Plant Drought Tolerance-Enhancing Bacterium. Baltrus DA, editor. Microbiol Resour Announc. 2018 Sep 27;7(12):e00842–18.

80. Yao J, Wang L, Zhang W, Liu M, Niu J. Effects of Bacillus megaterium on growth performance, serum biochemical parameters, antioxidant capacity, and immune function in suckling calves. Open Life Sci. 2020 Dec 31;15(1):1033–41.

81. Naumovich NI, Akhremchuk AE, Valentovich LN, Aleschenkov ZM, Ananyeva IN, Safronova GV. Molecular-genetic characterization of halotolerant strain *Priestia megaterium* BIM B-1314D. Dokl Akad nauk. 2022 Mar 9;66(1):55–64.

82. Ogawa K, Akagawa E, Yamane K, Sun ZW, LaCelle M, Zuber P, et al. The nasB operon and nasA gene are required for nitrate/nitrite assimilation in Bacillus subtilis. J Bacteriol. 1995 Mar;177(5):1409–13.

83. Walker V, Bruto M, Bellvert F, Bally R, Muller D, Prigent-Combaret C, et al. Unexpected Phytostimulatory Behavior for *Escherichia coli* and *Agrobacterium tumefaciens* Model Strains. MPMI. 2013 May;26(5):495–502.

84. Zeng Q, Wu X, Wang J, Ding X. Phosphate Solubilization and Gene Expression of Phosphate-Solubilizing Bacterium Burkholderia multivorans WS-FJ9 under Different Levels of Soluble Phosphate. Journal of Microbiology and Biotechnology. 2017;27(4):844–55.

85. Dai Z, Liu G, Chen H, Chen C, Wang J, Ai S, et al. Long-term nutrient inputs shift soil microbial functional profiles of phosphorus cycling in diverse agroecosystems. ISME J. 2020 Mar;14(3):757–70.

86. Ryu CM, Farag MA, Hu CH, Reddy MS, Wei HX, Paré PW, et al. Bacterial volatiles promote growth in *Arabidopsis*. Proc Natl Acad Sci USA. 2003 Apr 15;100(8):4927–32.

87. Bruto M, Prigent-Combaret C, Muller D, Moënne-Loccoz Y. Analysis of genes contributing to plant-beneficial functions in plant growth-promoting rhizobacteria and related Proteobacteria. Sci Rep. 2014 Sep 2;4:6261.

88. Petrova P, Velikova P, Petrov K. Genome Sequence of Bacillus velezensis 5RB, an Overproducer of 2,3-Butanediol. Dunning Hotopp JC, editor. Microbiol Resour Announc. 2019 Jan 3;8(1):e01475–18.

89. Spaepen S, Vanderleyden J. Auxin and plant-microbe interactions. Cold Spring Harb Perspect Biol. 2011 Apr 1;3(4):a001438.

90. Ma Q, Zhang X, Qu Y. Biodegradation and Biotransformation of Indole: Advances and Perspectives. Front Microbiol. 2018 Nov 1;9:2625.

91. Rai N, Kumar V, Sharma M, Akhter Y. Auxin transport mechanism of membrane transporter encoded by AEC gene of Bacillus licheniformis isolated from metagenome of Tapta Kund Hotspring of Uttrakhand, India. Int J Biol Macromol. 2021 Aug 31;185:277–86.

92. Maymon M, Martínez-Hidalgo P, Tran S, Ice T, Craemer K, Anbarchian T, et al. Mining the phytomicrobiome to understand how bacterial coinoculations enhance plant growth. Frontiers in Plant Science [Internet]. 2015 [cited 2022 Jul 10];6. Available from: https://www.frontiersin.org/articles/10.3389/fpls.2015.00784

93. Matteoli FP, Passarelli-Araujo H, Reis RJA, da Rocha LO, de Souza EM, Aravind L, et al. Genome sequencing and assessment of plant growth-promoting properties of a Serratia marcescens strain isolated from vermicompost. BMC Genomics. 2018 Oct 16;19(1):750.

94. Heylen K, Keltjens J. Redundancy and modularity in membrane-associated dissimilatory nitrate reduction in Bacillus. Frontiers in Microbiology. 2012;3:371.

95. Farhana A, Lappin SL. Biochemistry, Lactate Dehydrogenase. In: StatPearls [Internet]. Treasure Island (FL): StatPearls Publishing; 2022 [cited 2022 Jul 10]. Available from: http://www.ncbi.nlm.nih.gov/books/NBK557536/

96. Shin BS, Choi SK, Park SH. Regulation of the Bacillus subtilis Phosphotransacetylase Gene. Journal of Biochemistry. 1999 Aug 1;126(2):333– 9.

97. Choi SK, Saier MHJ. Mechanism of CcpA-Mediated Glucose Repression of the resABCDE Operon of Bacillus subtilis. MIP. 2006;11(1–2):104–10.

98. Duport C, Jobin M, Schmitt P. Adaptation in Bacillus cereus: From Stress to Disease. Frontiers in Microbiology [Internet]. 2016 [cited 2022 Jul 10];7. Available from: https://www.frontiersin.org/articles/10.3389/fmicb.2016.01550

99. Cruz Ramos H, Boursier L, Moszer I, Kunst F, Danchin A, Glaser P. Anaerobic transcription activation in Bacillus subtilis: identification of distinct FNR-dependent and -independent regulatory mechanisms. The EMBO Journal. 1995 Dec;14(23):5984–94.

100. Yin H, Li M, Lv M, Hepworth SR, Li D, Ma C, et al. SAUR15 Promotes Lateral and Adventitious Root Development via Activating H ^+^ -ATPases and Auxin Biosynthesis. Plant Physiol. 2020 Oct;184(2):837–51.

101. Sharma M, Singh D, Saksena HB, Sharma M, Tiwari A, Awasthi P, et al. Understanding the Intricate Web of Phytohormone Signalling in Modulating Root System Architecture. Int J Mol Sci. 2021 May 24;22(11):5508.

102. Zhao Y. Auxin Biosynthesis and Its Role in Plant Development. Annual Review of Plant Biology. 2010;61(1):49–64.

103. Bais HP, Weir TL, Perry LG, Gilroy S, Vivanco JM. The role of root exudates in rhizosphere interactions with plants and other organisms. Annu Rev Plant Biol. 2006;57:233–66.

104. Rudrappa T, Czymmek KJ, Paré PW, Bais HP. Root-Secreted Malic Acid Recruits Beneficial Soil Bacteria. Plant Physiology. 2008 Nov 6;148(3):1547–56.

105. Lo KJ, Lin SS, Lu CW, Kuo CH, Liu CT. Whole-genome sequencing and comparative analysis of two plant-associated strains of Rhodopseudomonas palustris (PS3 and YSC3). Sci Rep. 2018 Dec;8(1):12769.

106. Wang Y, Zhang W, Li K, Sun F, Han C, Wang Y, et al. Salt-induced plasticity of root hair development is caused by ion disequilibrium in Arabidopsis thaliana. J Plant Res. 2008 Jan;121(1):87–96.

107. Schleiff U. Why Root Morphology is Expected to Be a Key Factor for Crop Salt Tolerance? In: Pessarakli M, editor. Handbook of plant and crop stress. Fourth edition. Boca Raton: CRC Press, Taylor & Francis Group; 2020. p. 27.

108. Zhu JK. Abiotic Stress Signaling and Responses in Plants. Cell. 2016 Oct;167(2):313–24.

109. Yu P, Jiang N, Fu W, Zheng G, Li G, Feng B, et al. ATP Hydrolysis Determines Cold Tolerance by Regulating Available Energy for Glutathione Synthesis in Rice Seedling Plants. Rice (N Y). 2020 Apr 9;13(1):23.

110. Aslam S, Gul N, Mir MA, Asgher Mohd, Al-Sulami N, Abulfaraj AA, et al. Role of Jasmonates, Calcium, and Glutathione in Plants to Combat Abiotic Stresses Through Precise Signaling Cascade. Front Plant Sci. 2021 Jul 22;12:668029.

111. Gupta RS, Patel S, Saini N, Chen S. Robust demarcation of 17 distinct Bacillus species clades, proposed as novel Bacillaceae genera, by phylogenomics and comparative genomic analyses: description of Robertmurraya kyonggiensis sp. nov. and proposal for an emended genus Bacillus limiting it only to the members of the Subtilis and Cereus clades of species. International Journal of Systematic and Evolutionary Microbiology. 2020 Nov 1;70(11):5753–98.

112. International Code of Nomenclature of Prokaryotes. Int J Syst Evol Microbiol. 2019 Jan;69(1A):S1–111.

113. Shivaji S, Chaturvedi P, Begum Z, Pindi PK, Manorama R, Padmanaban DA, et al. Janibacter hoylei sp. nov., Bacillus isronensis sp. nov. and Bacillus aryabhattai sp. nov., isolated from cryotubes used for collecting air from the upper atmosphere. INTERNATIONAL JOURNAL OF SYSTEMATIC AND EVOLUTIONARY MICROBIOLOGY. 2009 Dec 1;59(12):2977–86.

114. Bary A. Vergleichende Morphologie und Biologie der Pilze, Mycetozoen und Bacterien. Leipzig, Germany: Engelmann; 1884. 586 p.

115. Narsing Rao MP, Dong ZY, Liu GH, Li L, Xiao M, Li WJ. Reclassification of Bacillus aryabhattai Shivaji et al. 2009 as a later heterotypic synonym of Bacillus megaterium de Bary 1884 (Approved Lists 1980). FEMS Microbiol Lett. 2019 Nov 1;366(22):fnz258.

116. Silva P, Gerós H. Regulation by salt of vacuolar H ^+^ -ATPase and H ^+^ - pyrophosphatase activities and Na ^+^ /H ^+^ exchange. Plant Signaling & Behavior. 2009 Aug;4(8):718–26.

117. Ahmad P, Azooz MM, Prasad MNV, editors. Ecophysiology and Responses of Plants under Salt Stress [Internet]. NY, USA: Springer New York; 2013 [cited 2022 Jul 10]. Available from: http://link.springer.com/10.1007/978-1-4614-4747-4

118. Sellami S, Le Hir R, Thorpe MR, Vilaine F, Wolff N, Brini F, et al. Salinity Effects on Sugar Homeostasis and Vascular Anatomy in the Stem of the Arabidopsis Thaliana Inflorescence. IJMS. 2019 Jun 28;20(13):3167.

119. Sasse J, Martinoia E, Northen T. Feed Your Friends: Do Plant Exudates Shape the Root Microbiome? Trends in Plant Science. 2018 Jan;23(1):25–41.

120. Trivedi P, Leach JE, Tringe SG, Sa T, Singh BK. Plant–microbiome interactions: from community assembly to plant health. Nat Rev Microbiol. 2020 Nov;18(11):607–21.

121. Nobori T, Velásquez AC, Wu J, Kvitko BH, Kremer JM, Wang Y, et al. Transcriptome landscape of a bacterial pathogen under plant immunity. Proc Natl Acad Sci USA. 2018 Mar 27;115(13):E3055–64.

122. Nobori T, Cao Y, Entila F, Dahms E, Tsuda Y, Garrido-Oter R, et al. Dissecting the cotranscriptome landscape of plants and their microbiota. EMBO reports. 2022 Dec 6;23(12):e55380.

123. Nobori T, Wang Y, Wu J, Stolze SC, Tsuda Y, Finkemeier I, et al. Multidimensional gene regulatory landscape of a bacterial pathogen in plants. Nat Plants. 2020 Jul;6(7):883–96.

124. Leake JR, Ostle NJ, Rangel-Castro JI, Johnson D. Carbon fluxes from plants through soil organisms determined by field 13CO2 pulse-labelling in an upland grassland. Applied Soil Ecology. 2006 Sep;33(2):152–75.

125. Hacquard S, Spaepen S, Garrido-Oter R, Schulze-Lefert P. Interplay Between Innate Immunity and the Plant Microbiota. Annu Rev Phytopathol. 2017 Aug 4;55(1):565–89.

126. Khare E, Mishra J, Arora NK. Multifaceted interactions between endophytes and plant: developments and prospects. Front Microbiol. 2018 Nov 15;9:2732.

