## Supplementary material for "A cyclic dipeptide for salinity stress alleviation and the trophic flexibility of an endophyte reveal niches in salt marsh plant-microbe interactions": priest01_sup_fig.BioRxiv.pdf

**Article title:**

**Affiliations:**

### Supplementary Figures

**Fig. S1** Volcano plot of differential expression genes (DEGs).

**Fig. S2** Gene set enrichment analysis (GSEA) of DEGs (BP01R2 vs CK) within biological process GO ontologies.

**Fig. S3** Gene set enrichment analysis (GSEA) of DEGs (BP01R2 vs CK) within molecular functions GO ontologies.

**Fig. S4** Gene set enrichment analysis (GSEA) of DEGs (BP01R2 vs CK) within KEGG pathways.

**Fig. S5** Expression pattern of DEGs (BP01R2 vs CK) in KEGG glycolysis and gluconeogenesis pathway.

**Fig. S6** Expression pattern of DEGs (BP01R2 vs CK) in KEGG oxidative phosphorylation pathway.

**Fig. S7** Expression pattern of DEGs (BP01R2 vs CK) in KEGG plant hormone signal transduction pathway.

**Fig. S8** Gene set enrichment analysis (GSEA) of DEGs (BP01R2_NaCl vs CK_NaCl) within biological process GO ontologies.

**Fig. S9** Gene set enrichment analysis (GSEA) of DEGs (BP01R2_NaCl vs CK_NaCl) within KEGG pathways.

**Fig. S10** Expression pattern of DEGs (BP01R2_NaCl vs CK_NaCl) in KEGG plant hormone signal transduction pathway.

**Fig. S11** Expression pattern of DEGs (BP01R2_NaCl vs CK_NaCl) in MAPK signaling pathway.

**Fig. S12** Expression pattern of DEGs (BP01R2_NaCl vs CK_NaCl) in ribosome pathway.

**Fig. S13** Expression pattern of DEGs (BP01R2_NaCl vs CK_NaCl) in KEGG glutathione metabolism pathway.

**Fig. S14** Expression pattern of DEGs (CK_NaCl compared CK) in KEGG plant hormone signal transduction pathway.

**Fig. S15** Expression pattern of DEGs (CK_NaCl compared CK) in KEGG MAPK signaling pathway.

**Fig. S16** Expression pattern of DEGs (CK_NaCl compared CK) in KEGG ribosome pathway.

**Fig. S17** Expression pattern of DEGs (CK_NaCl compared CK) in glutathione metabolism pathway.

**Fig. S18** IAA production evaluation of BP01R2.

**Fig. S19** Genome map of *Priestia megaterium* BP01R2 chromosome.

**Fig. S20** ABC transporters identified within the genome of BP01R2.

**Fig. S21** Two-component systems identified within the genome of BP01R2.

**Fig. S22** The multiple sequence alignment analysis of oligoribonuclease NrnB among *Priestia megaterium* and its relatives.

**Supplementary Figures**

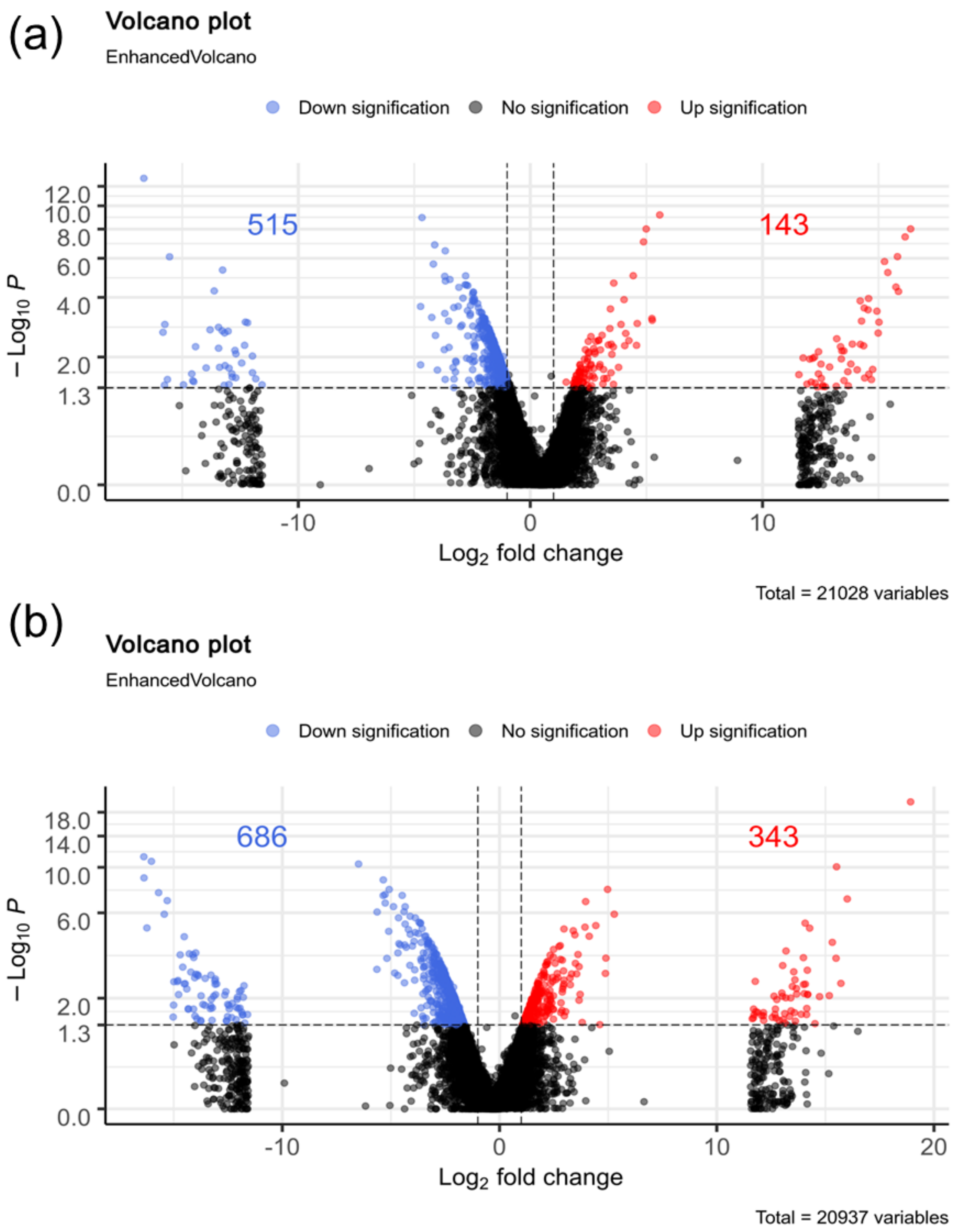


**Fig. S1 Volcano plot of differential expression genes (DEGs). (a)** BP01R2 vs CK. **(b)** BP01R2_NaCl vs CK_NaCl.

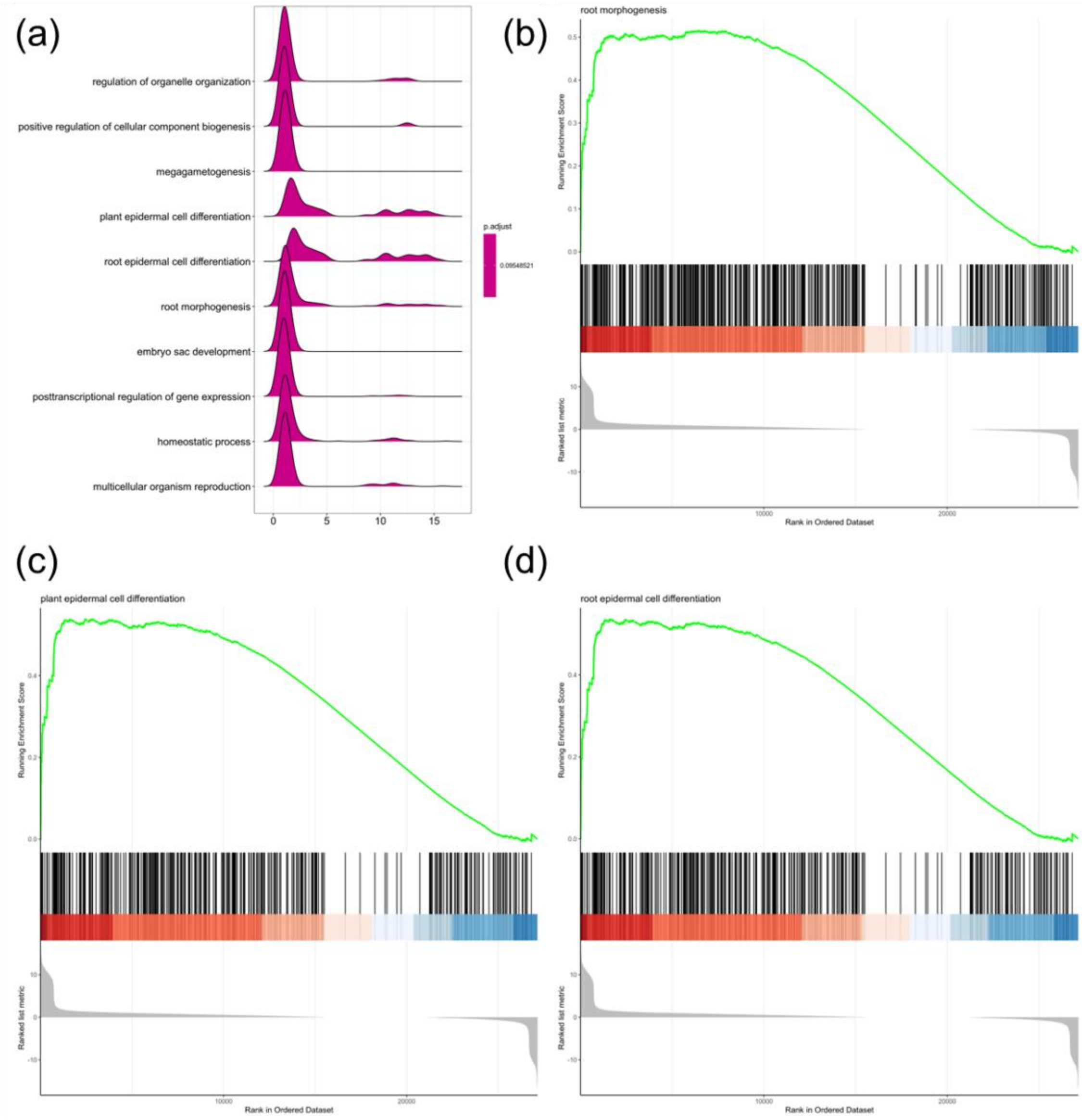


**Fig. S2 Gene set enrichment analysis (GSEA) of DEGs (BP01R2 vs CK) within biological process GO ontologies. (a)** Ridgeline plot for expression distribution. GSEA score plots of **(b)** root morphogenesis, **(c)** plant epidermal cell differentiation and **(d)** root epidermal cell differentiation.

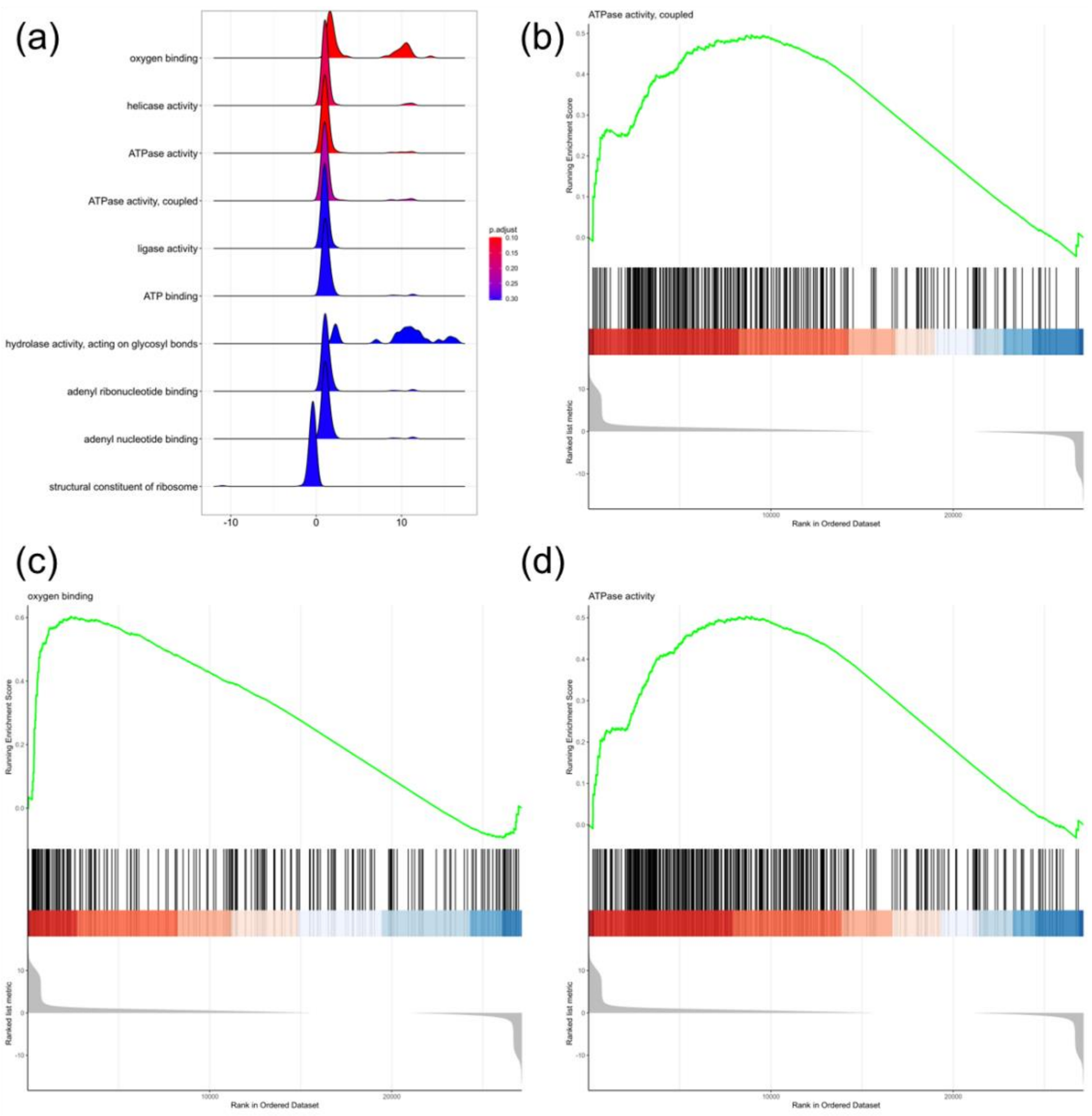


**Fig. S3 Gene set enrichment analysis (GSEA) of DEGs (BP01R2 vs CK) within molecular functions GO ontologies. (a)** Ridgeline plot for expression distribution. GSEA score plots of **(b)** ATPase activity, coupled, **(c)** oxygen binding and **(d)** ATPase activity.

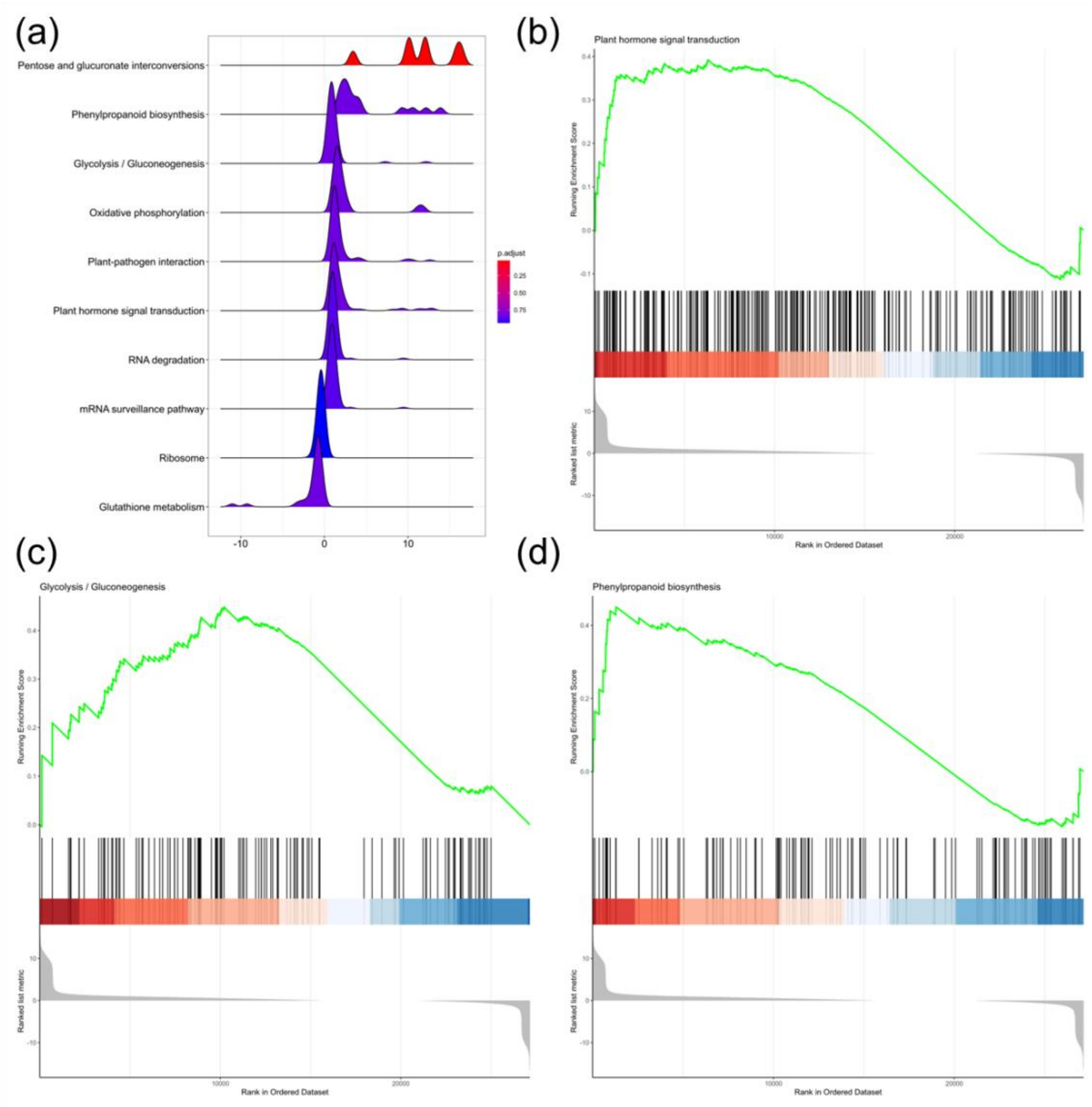


**Fig. S4 Gene set enrichment analysis (GSEA) of DEGs (BP01R2 vs CK) within KEGG pathways. (a)** Ridgeline plot for expression distribution. GSEA score plots of **(b)** plant hormone signal transduction, **(c)** glycolysis/gluconeogenesis and **(d)** phenylpropanoid biosynthesis.

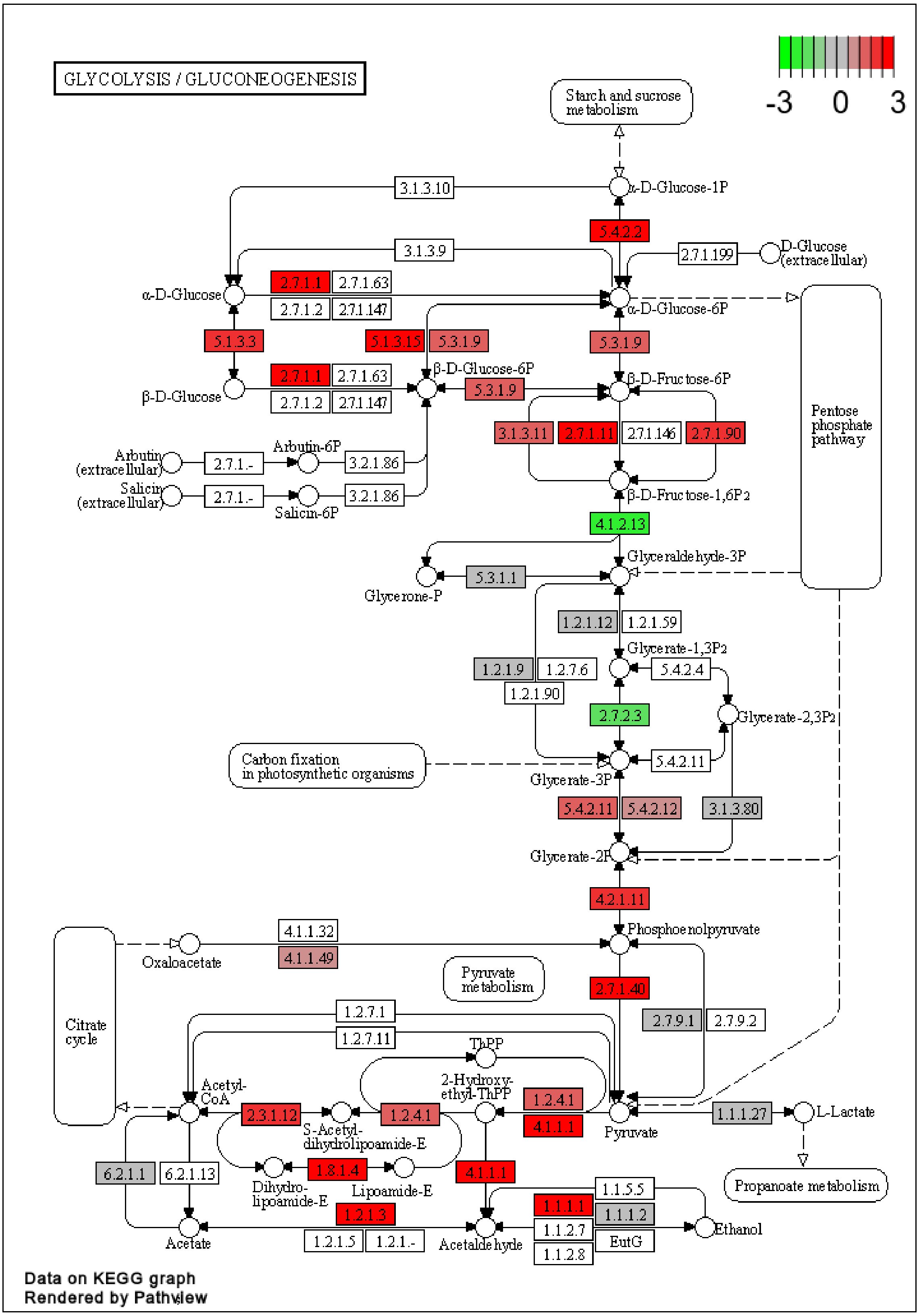


**Fig. S5 Expression pattern of DEGs (BP01R2 vs CK) in KEGG glycolysis and gluconeogenesis pathway.**

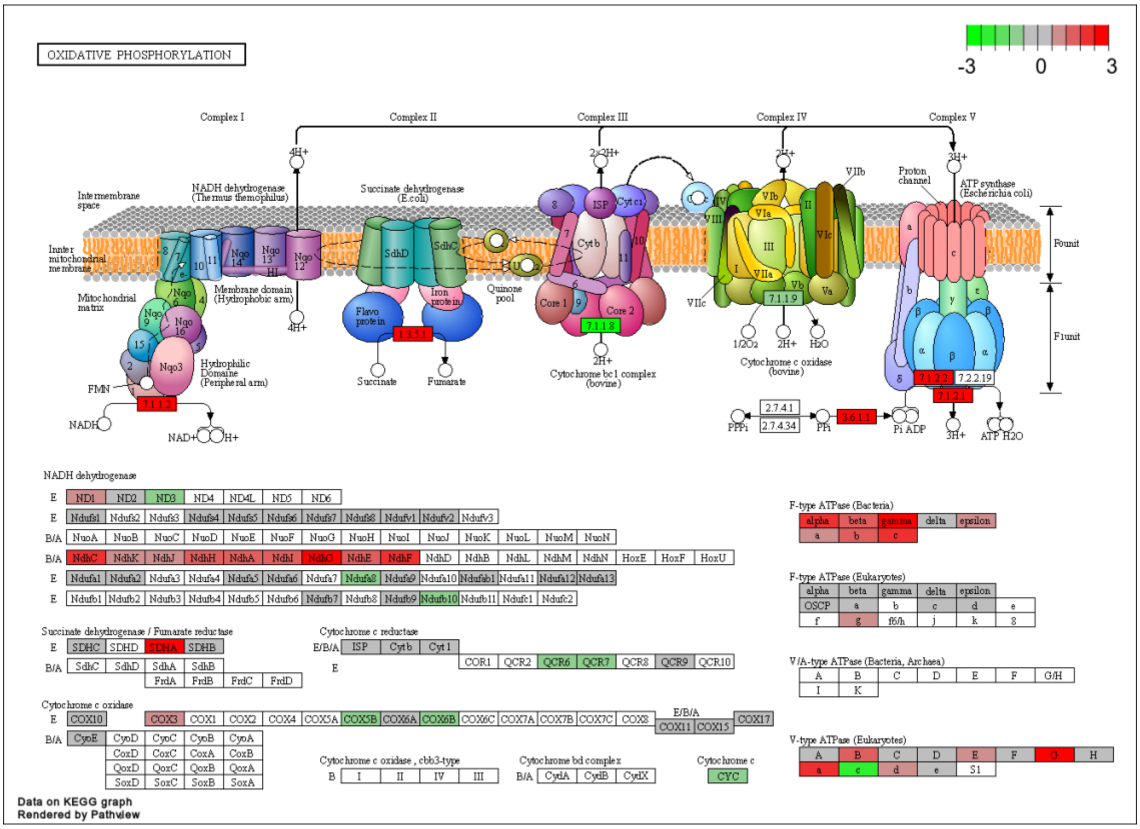


**Fig. S6 Expression pattern of DEGs (BP01R2 vs CK) in KEGG oxidative phosphorylation pathway.**

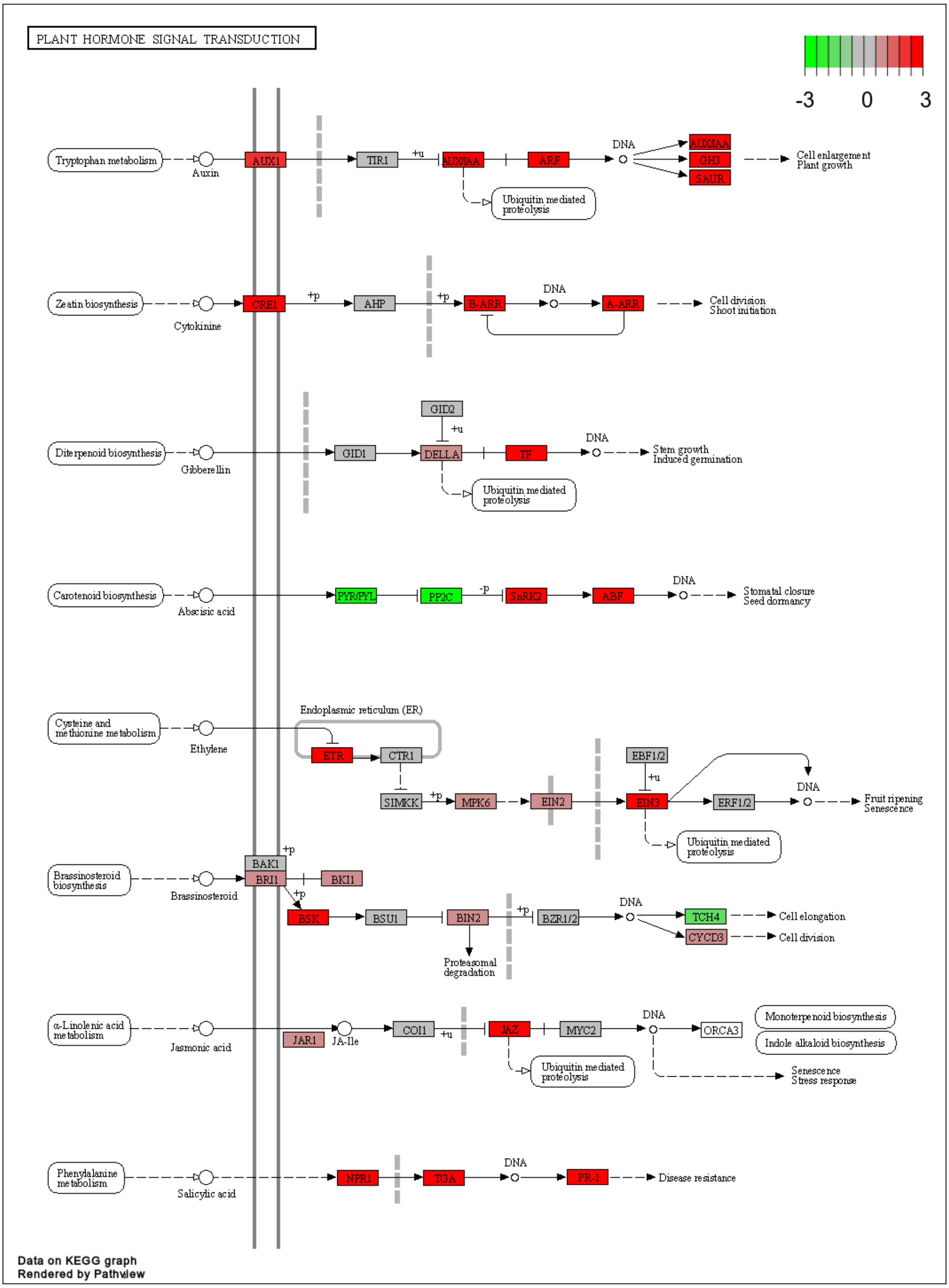


**Fig. S7 Expression pattern of DEGs (BP01R2 vs CK) in KEGG plant hormone signal transduction pathway.**

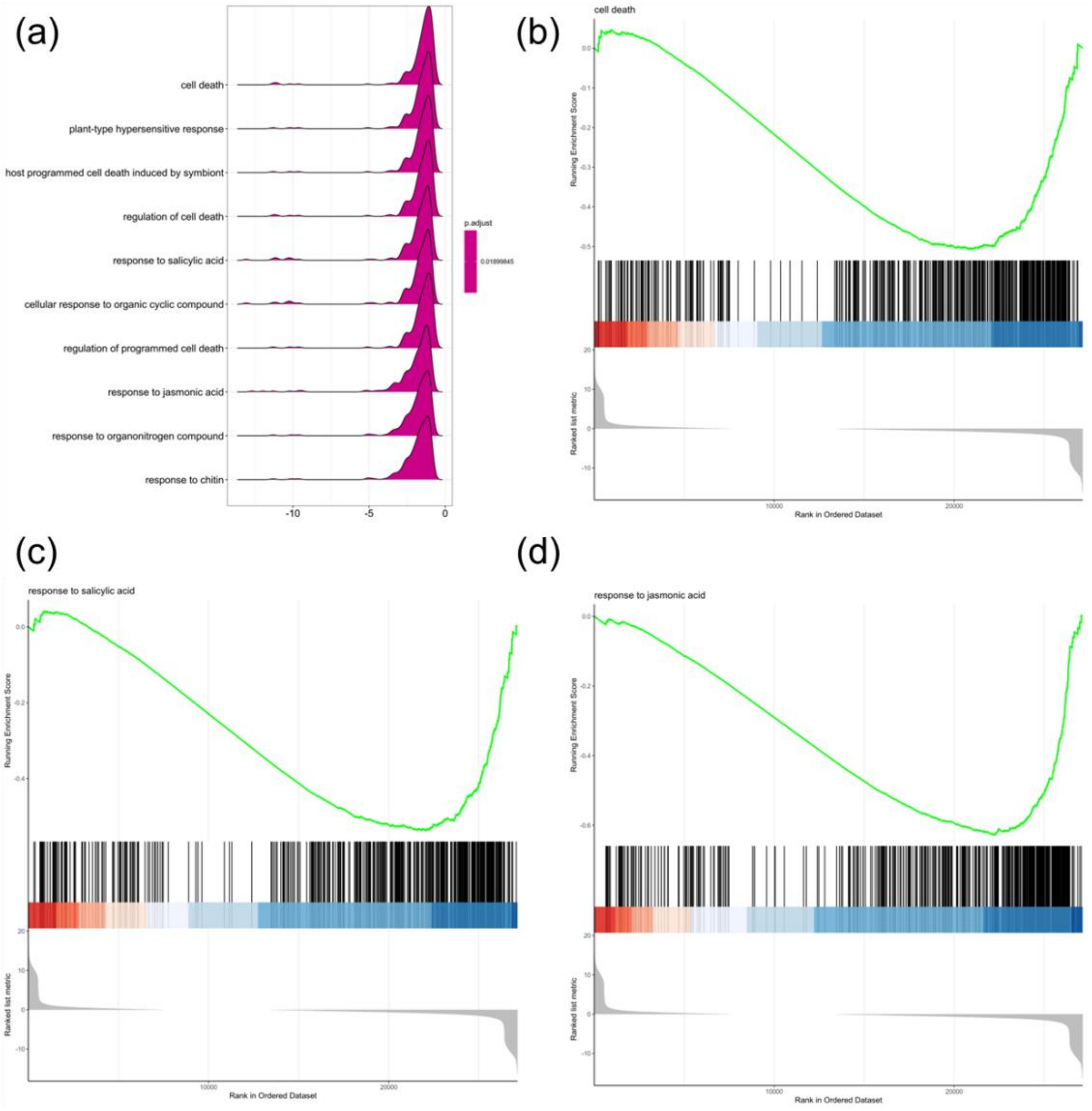


**Fig. S8 Gene set enrichment analysis (GSEA) of DEGs (BP01R2_NaCl vs CK_NaCl) within biological process GO ontologies. (a)** Ridgeline plot for expression distribution. GSEA score plots of **(b)** cell death, **(c)** response to salicylic acid and **(d)** response to jasmonic acid.

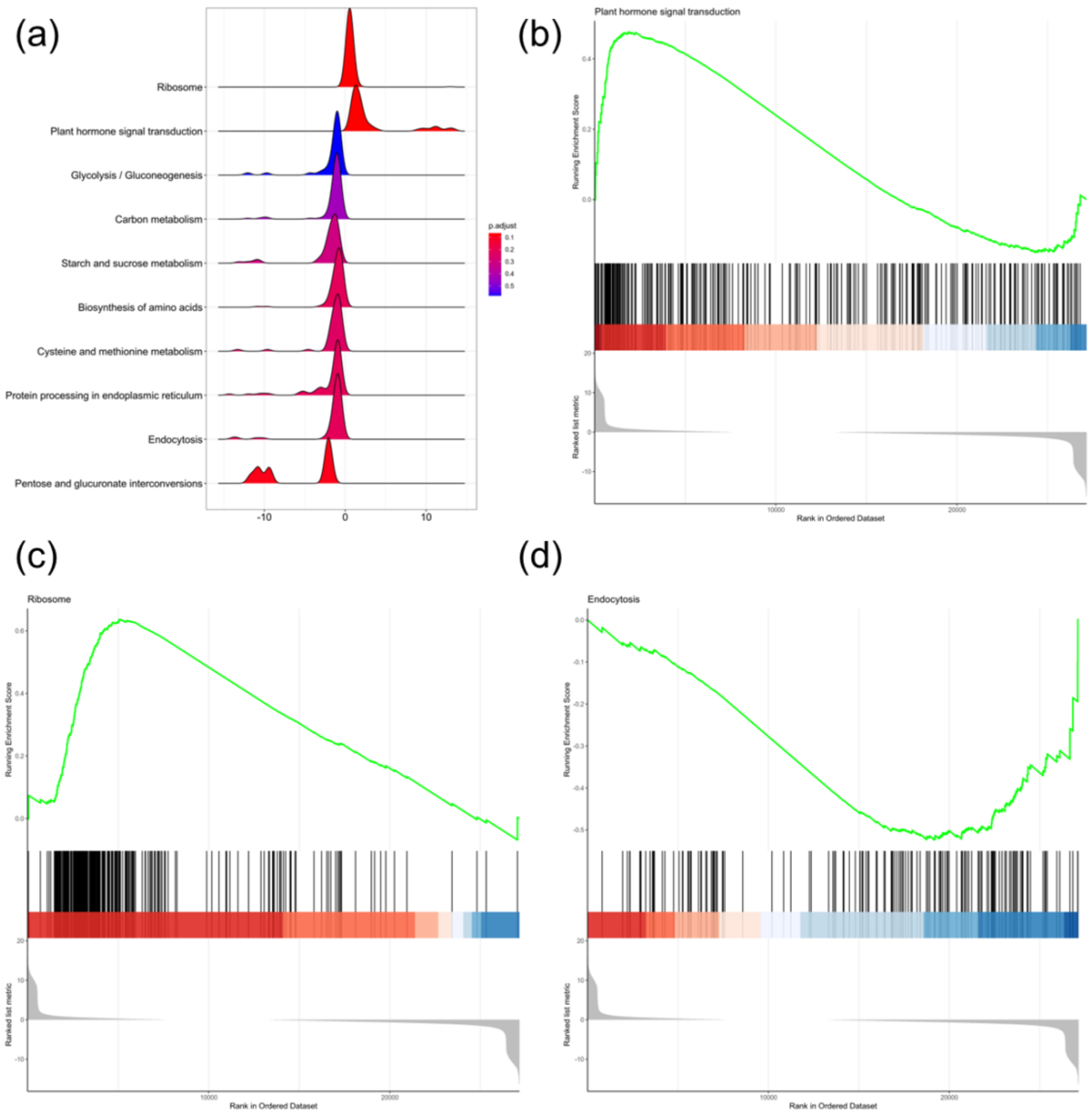


**Fig. S9 Gene set enrichment analysis (GSEA) of DEGs (BP01R2_NaCl vs CK_NaCl) within KEGG pathways. (a)** Ridgeline plot for expression distribution. GSEA score plots of **(b)** plant hormone signal transduction, **(c)** ribosome and **(d)** endocytosis.

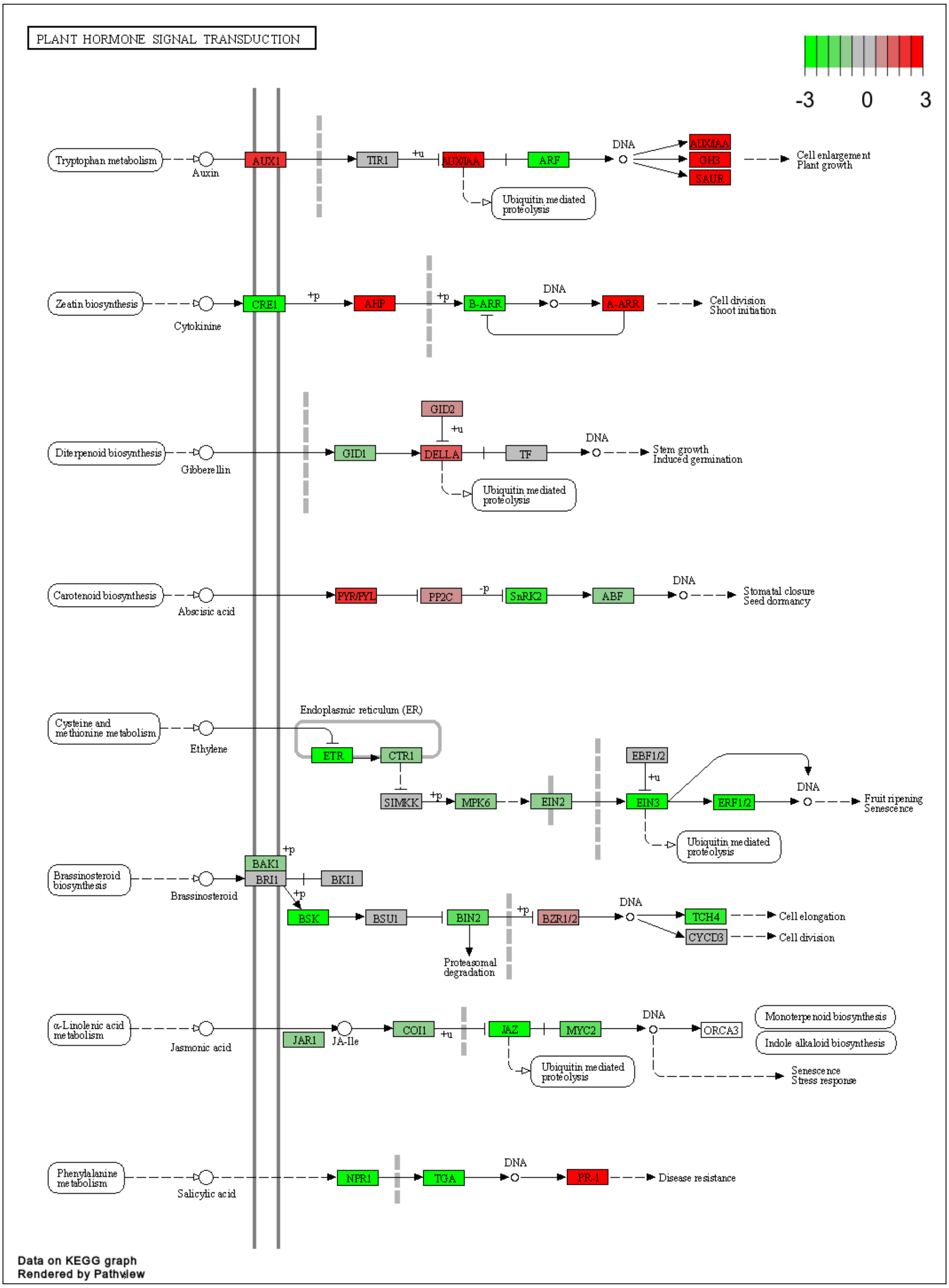


**Fig. S10 Expression pattern of DEGs (BP01R2_NaCl vs CK_NaCl) in KEGG plant hormone signal transduction pathway.**

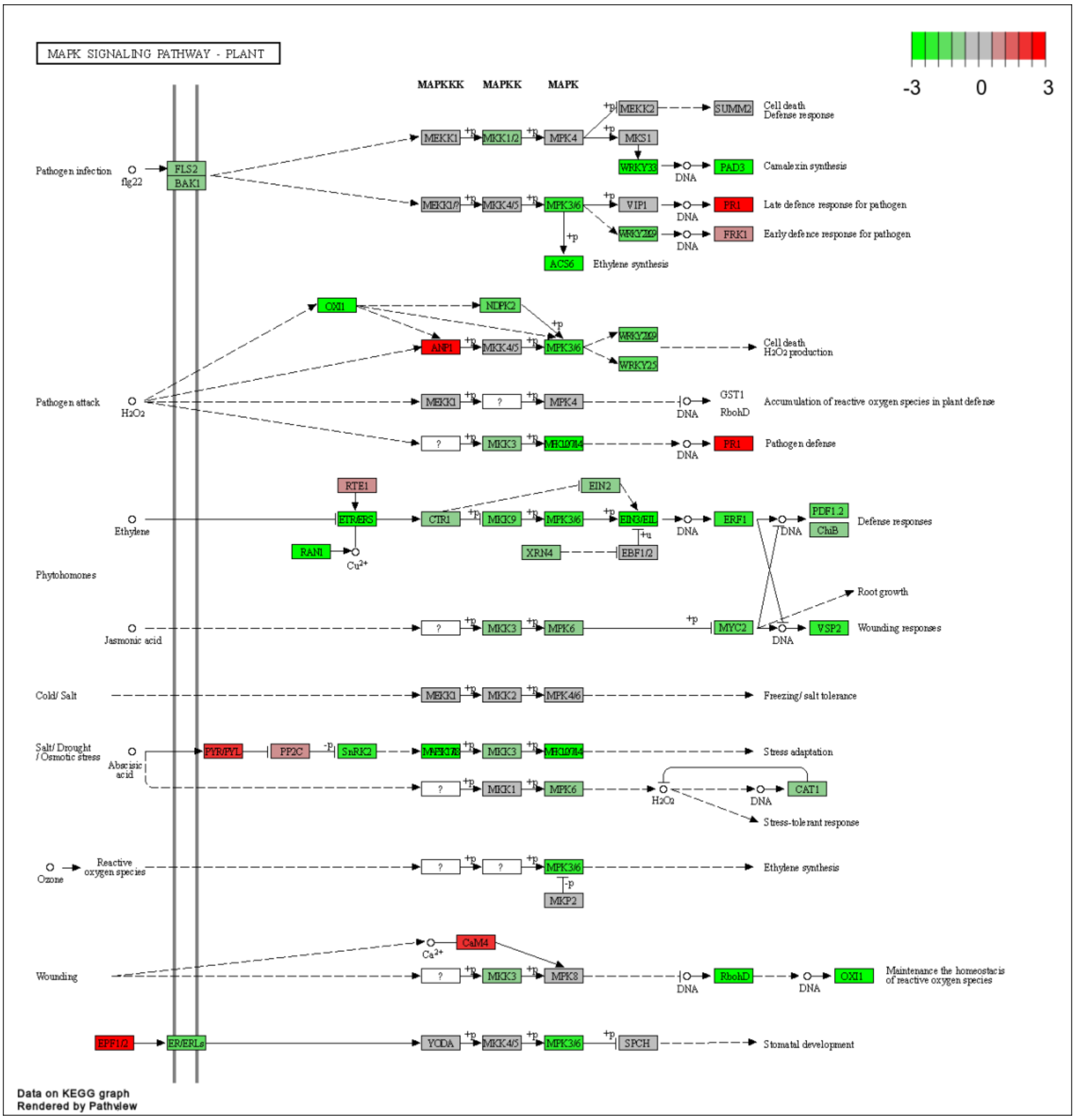


**Fig. S11 Expression pattern of DEGs (BP01R2_NaCl vs CK_NaCl) in MAPK signaling pathway.**

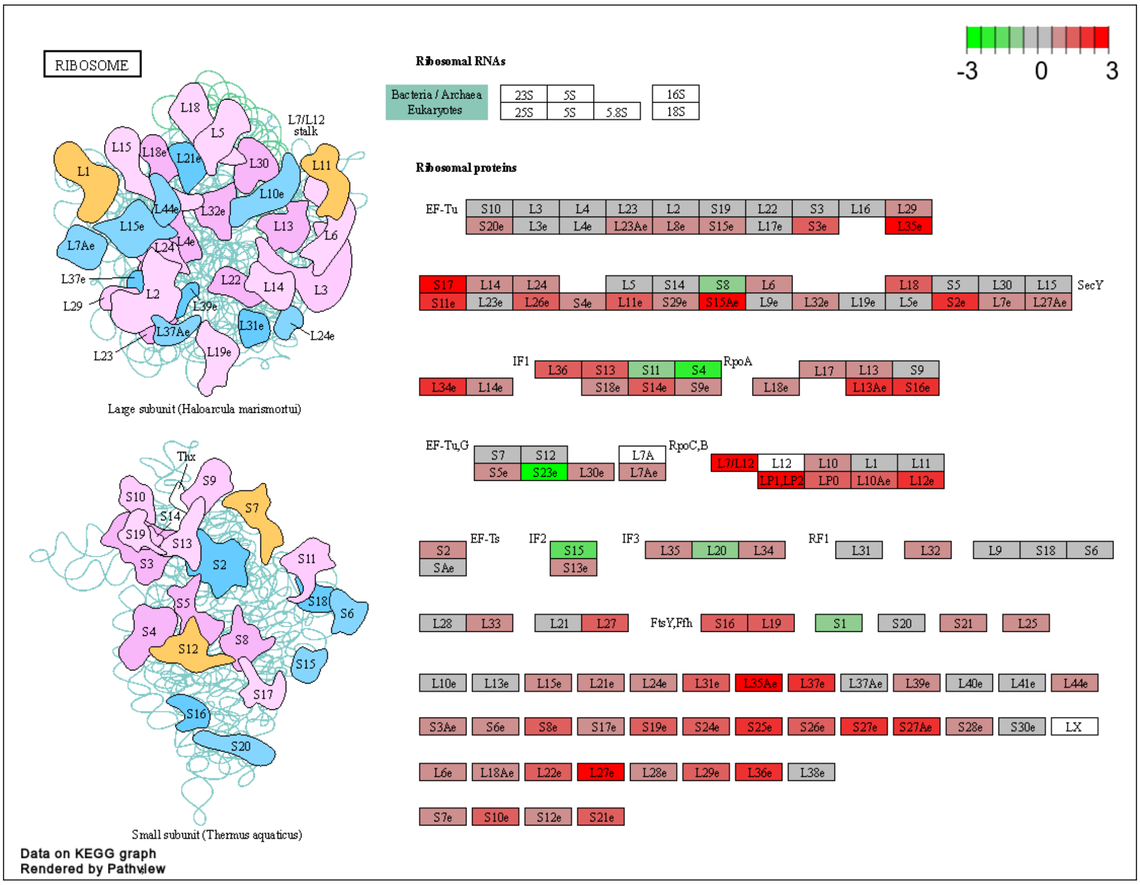


**Fig. S12 Expression pattern of DEGs (BP01R2_NaCl vs CK_NaCl) in ribosome pathway.**

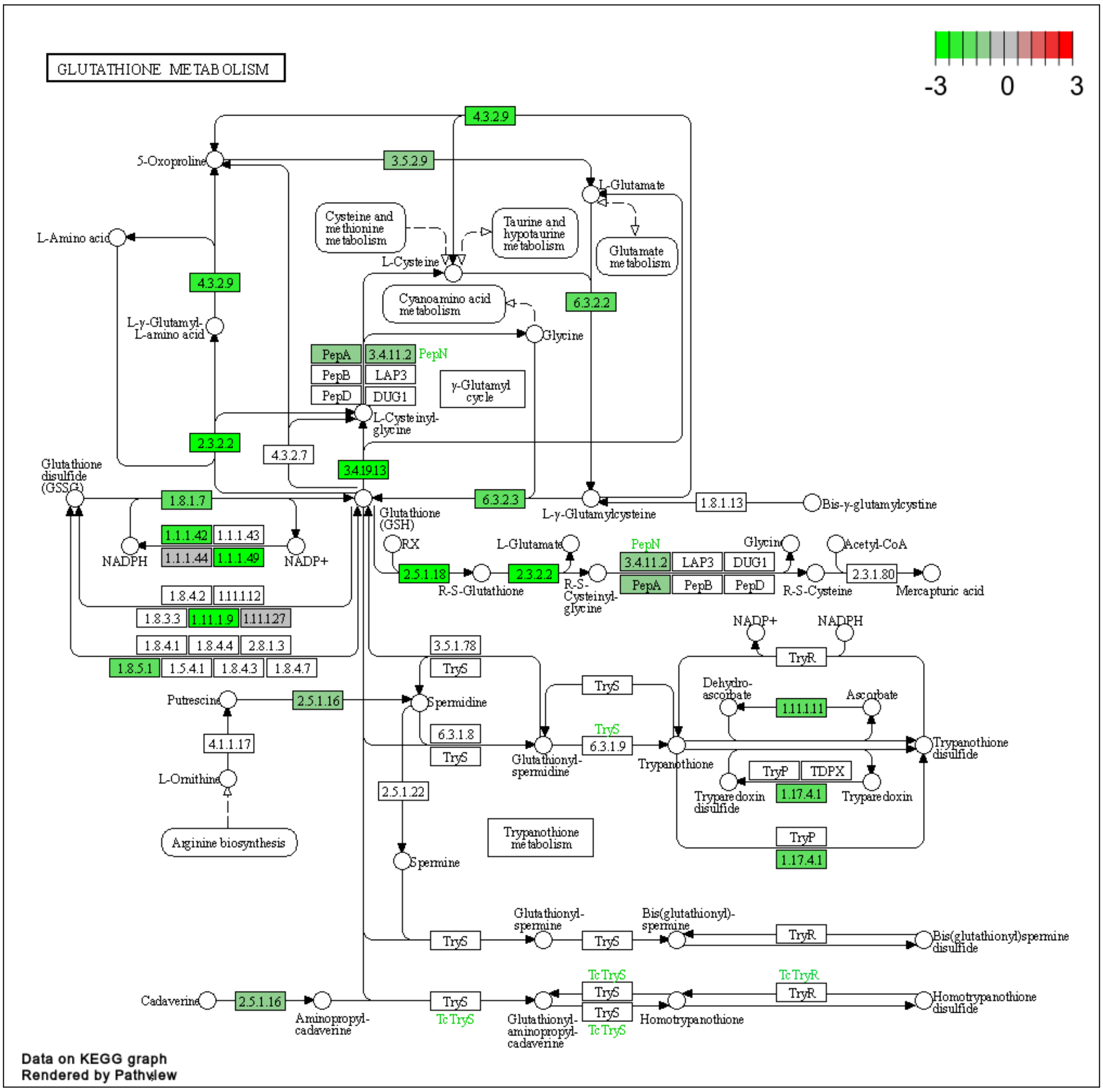


**Fig. S13 Expression pattern of DEGs (BP01R2_NaCl vs CK_NaCl) in KEGG glutathione metabolism pathway.**

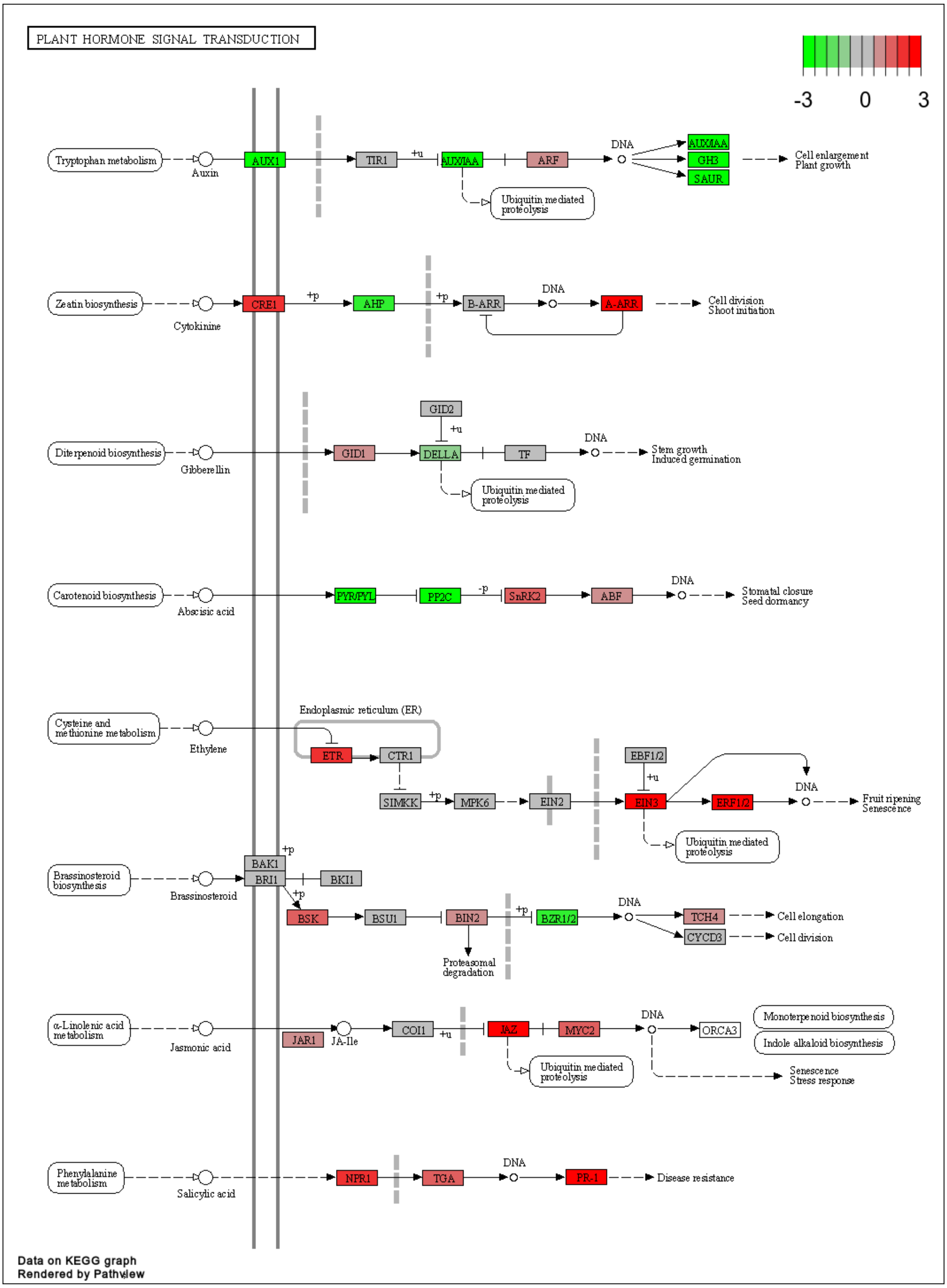


**Fig. S14 Expression pattern of DEGs (CK_NaCl compared CK) in KEGG plant hormone signal transduction pathway.**

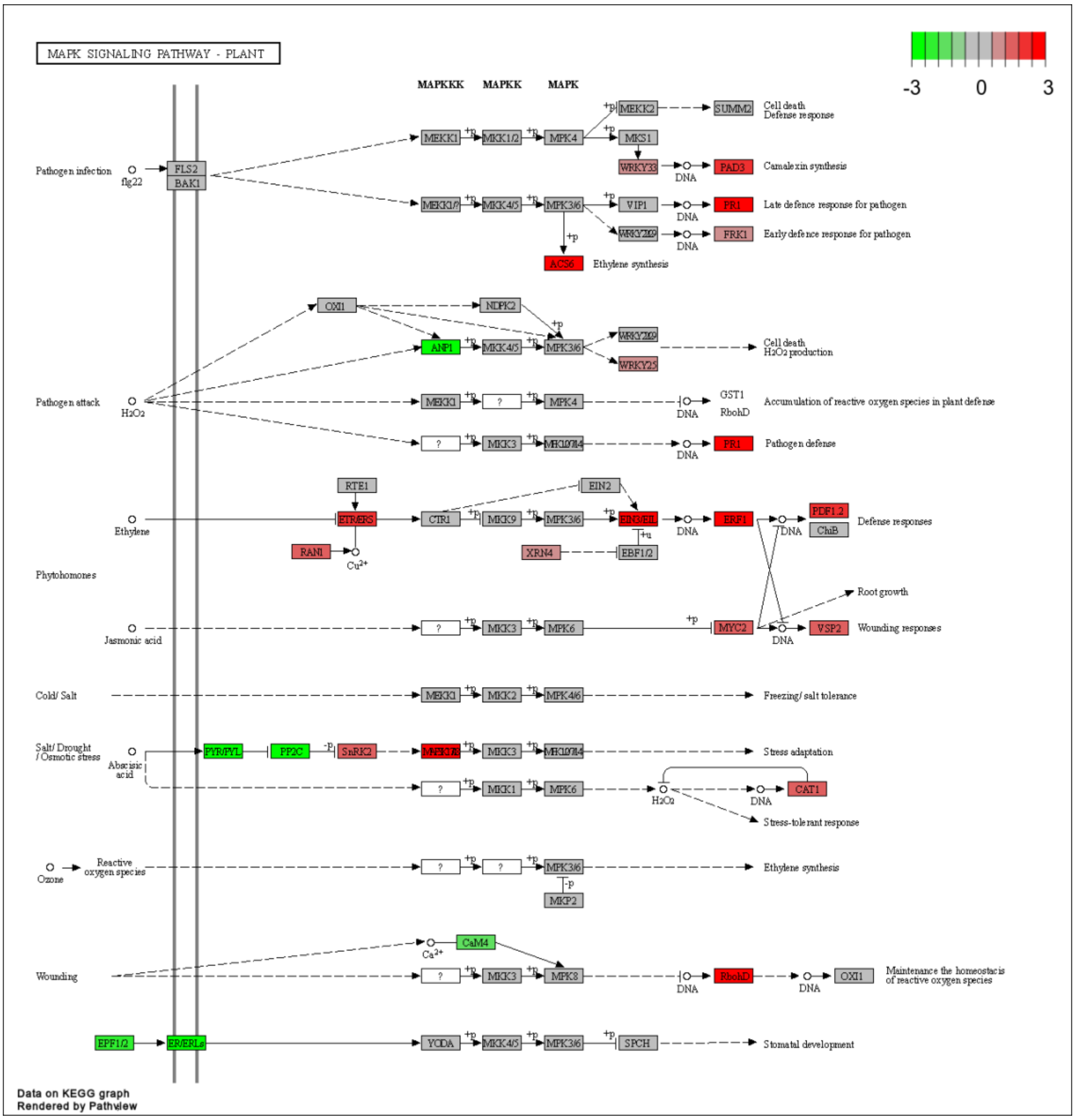


**Fig. S15 Expression pattern of DEGs (CK_NaCl compared CK) in KEGG MAPK signaling pathway.**

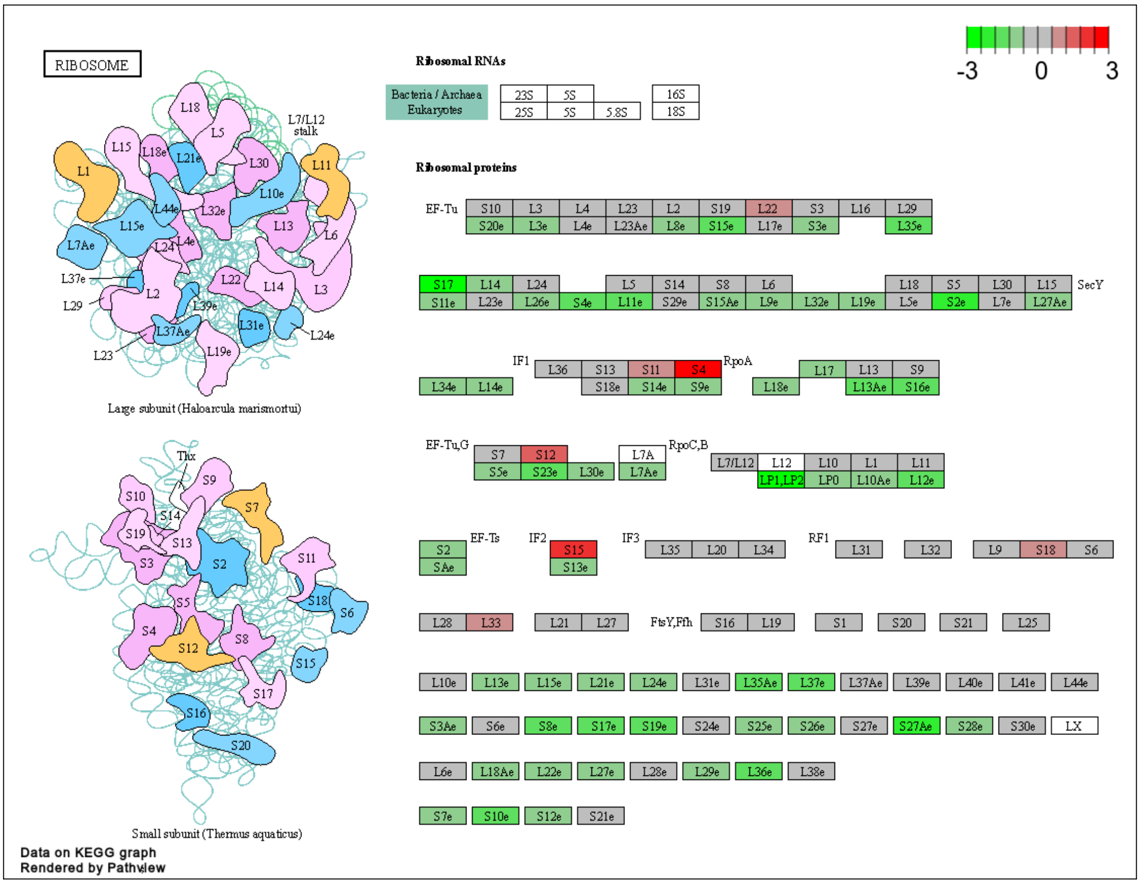


**Fig. S16 Expression pattern of DEGs (CK_NaCl compared CK) in KEGG ribosome pathway.**

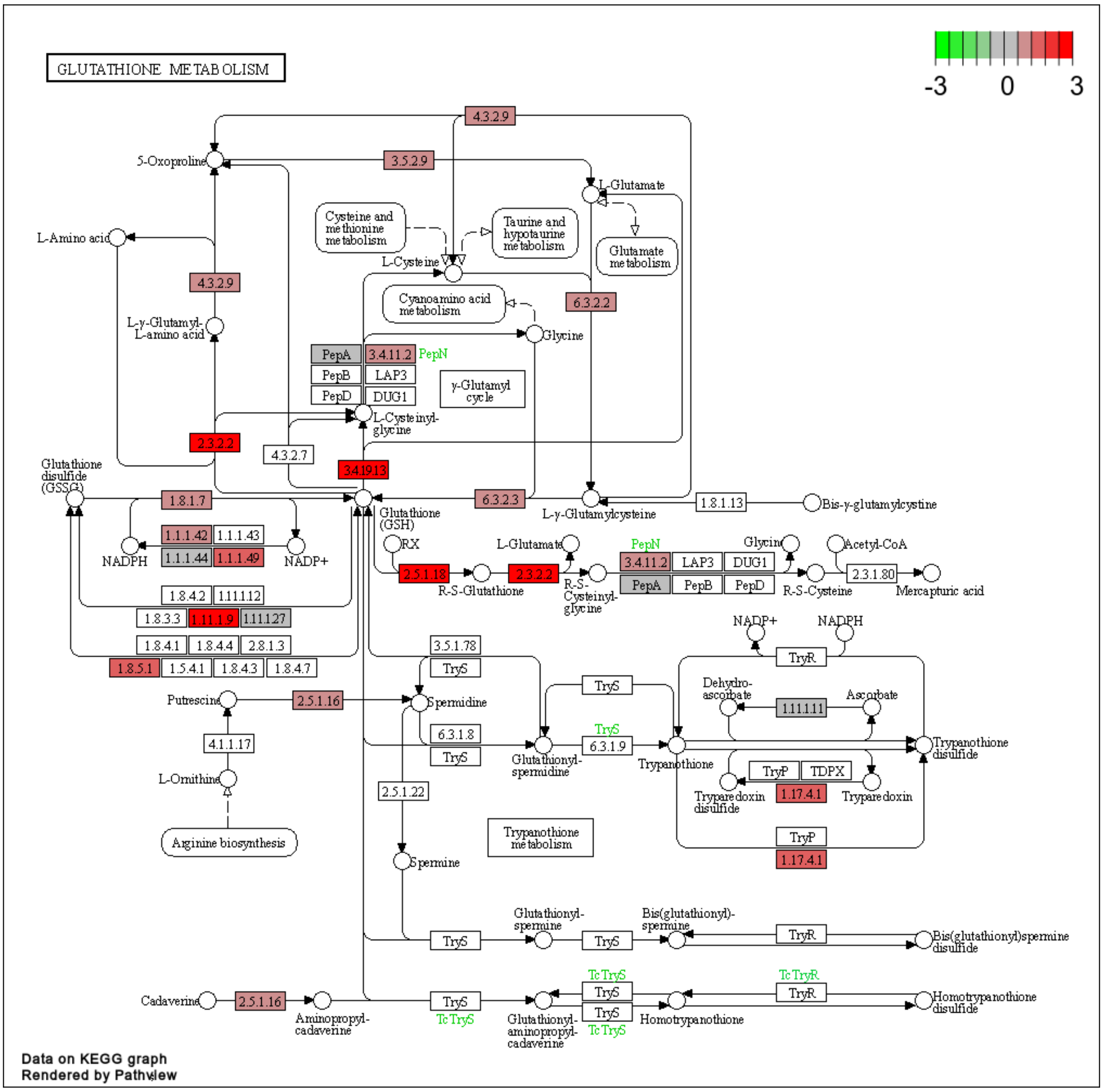


**Fig. S17 Expression pattern of DEGs (CK_NaCl compared CK) in glutathione metabolism pathway.**

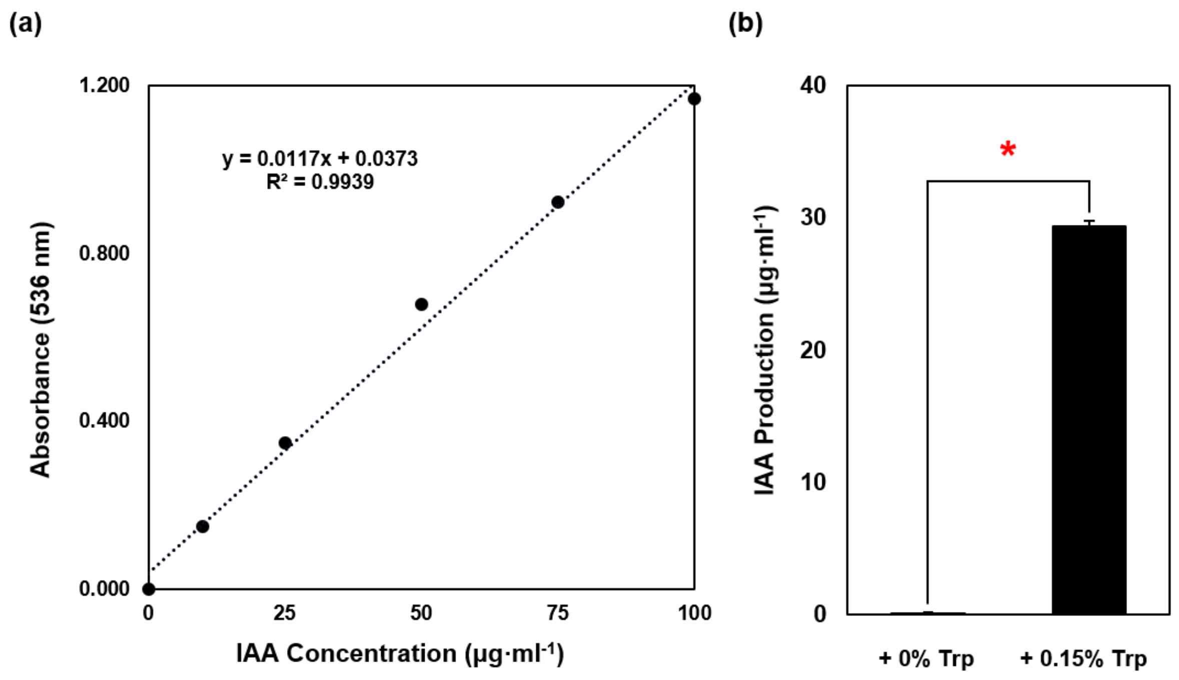


**Fig. S18 IAA production evaluation of BP01R2. (a)** Calibration curve of IAA. **(b)** IAA production was measured by Salkowski reagent methods after 24 hours of incubation with or without 0.15% L-Tryptophan (Trp) supplementation. Each sample was performed with at least three biological and three technical replications. *$p<0.05$.

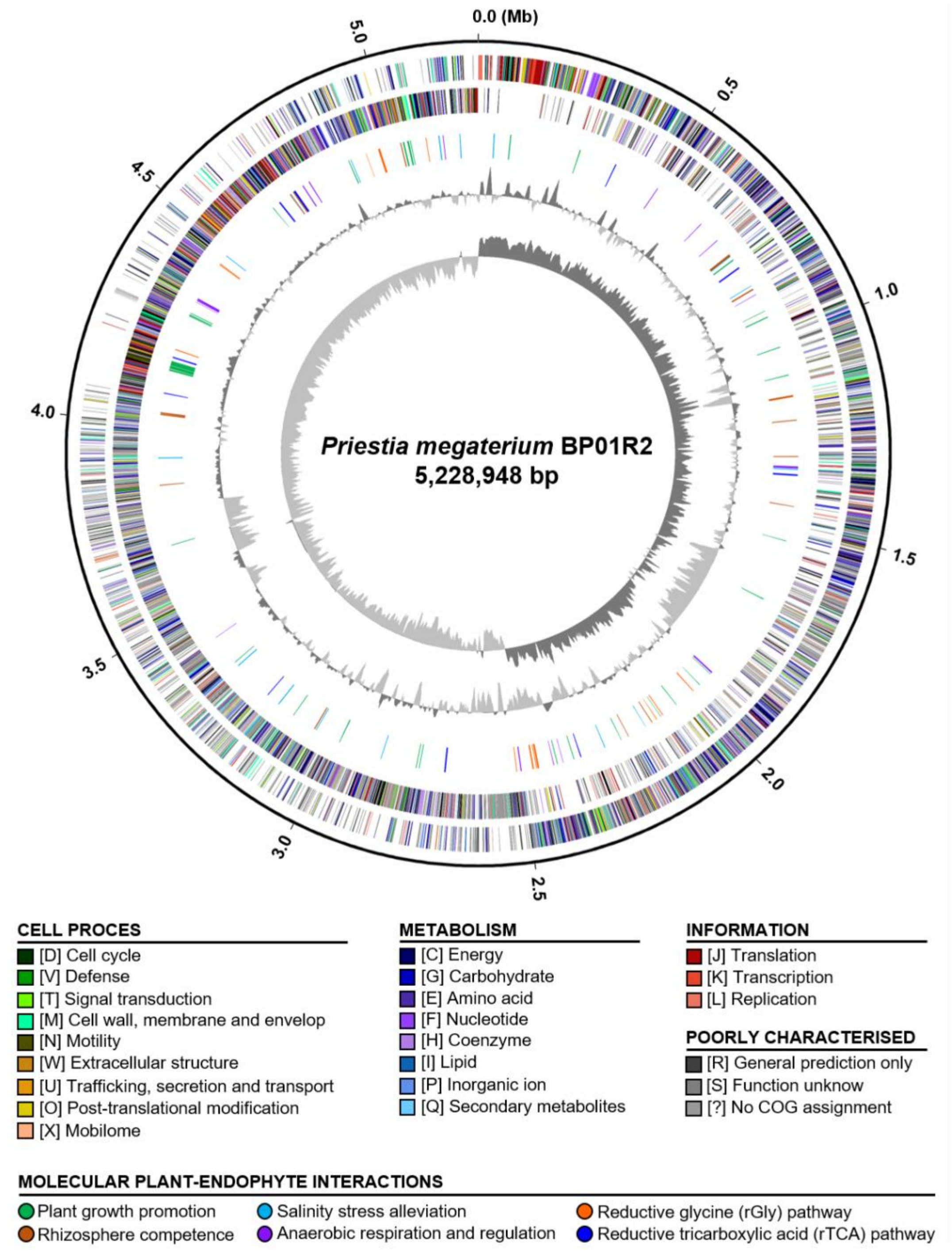


**Fig. S19 Genome map of *Priestia megaterium* BP01R2 chromosome.** Rings from outside in: (1) Scale marks (Mb). (2 and 3) Forward and reverse strand of coding sequences, respectively. Colour-coded by functional categories (square). (4) Genes associated with identical molecular plant-endophyte interactions. Colour-coded by identified genes involved in metabolism and pathway (circle). See the gene name and locus accession details in Table S6. (5) GC content (dark grey, above average; light grey, average). (6) GC skew (dark grey, positive; light grey, negative).

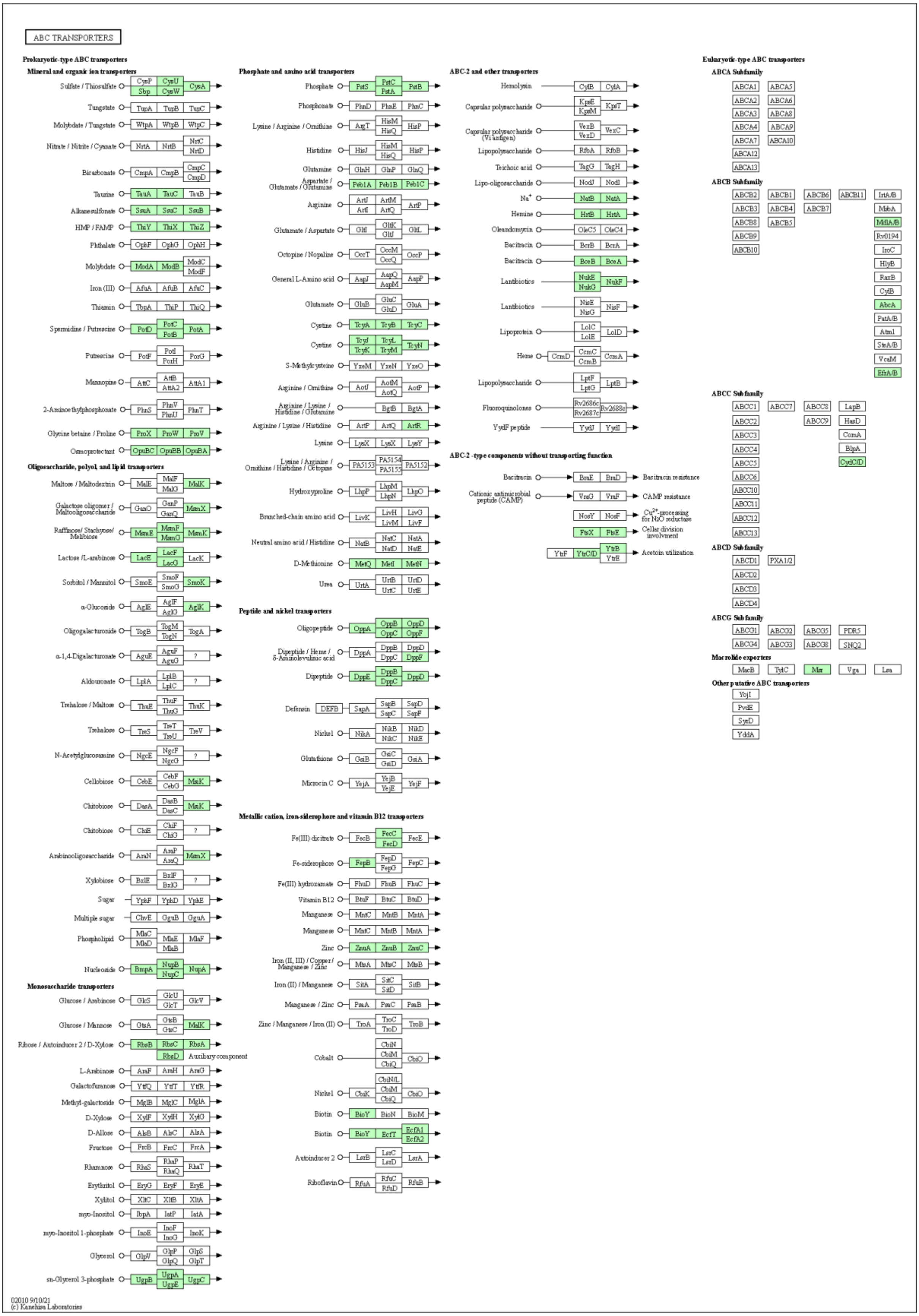


**Fig. S20 ABC transporters identified within the genome of BP01R2.** The green coloured genes are present in the BP01R2 genome.

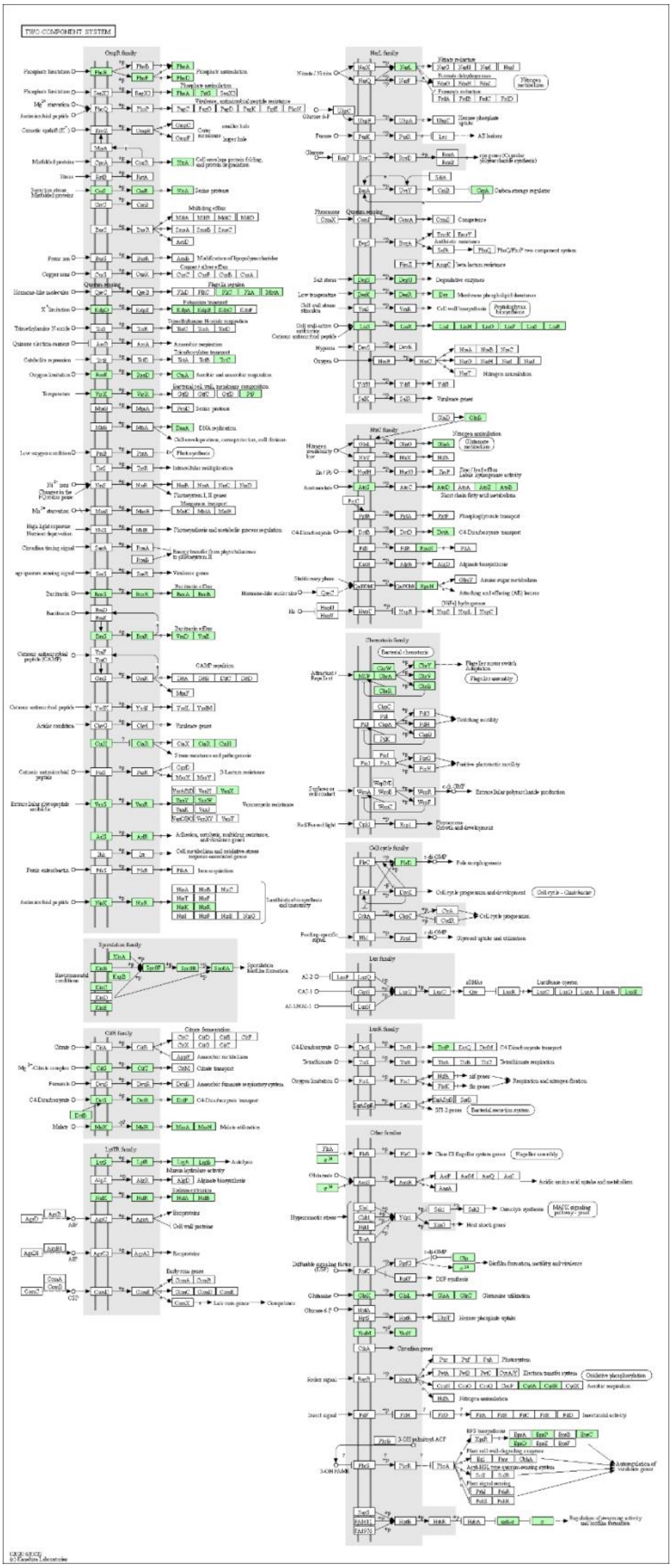


**Fig. S21 Two-component systems identified within the genome of BP01R2.** The green coloured genes are present in the BP01R2 genome.

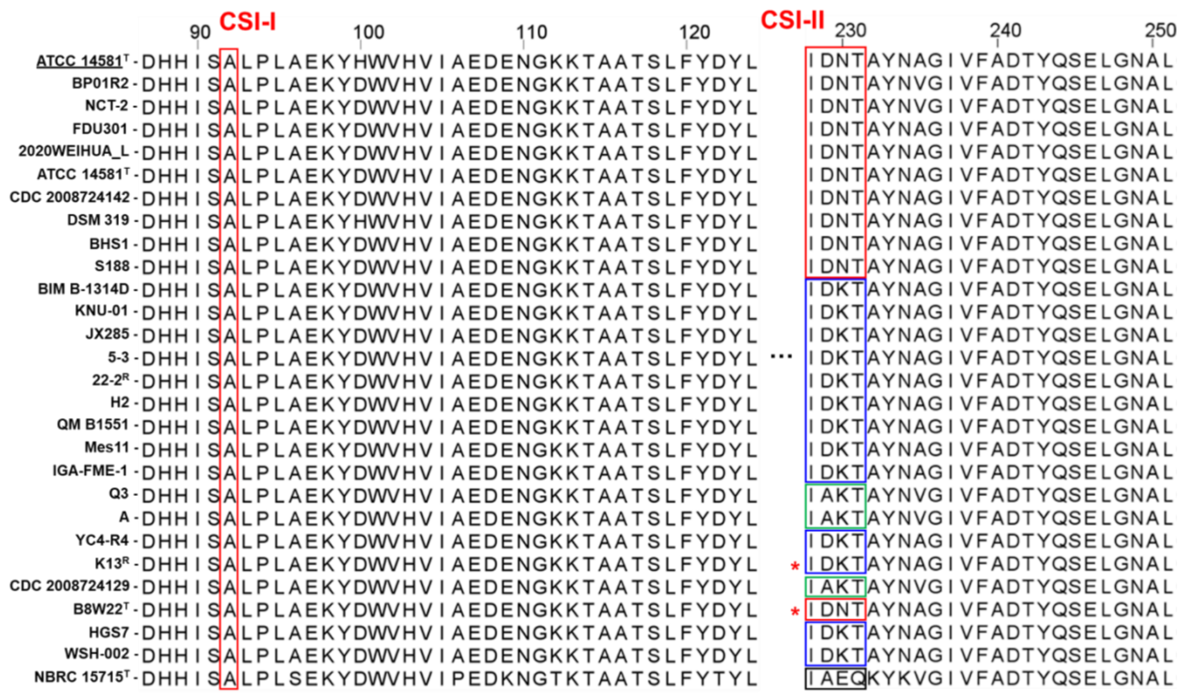

**Fig. S22 The multiple sequence alignment analysis of oligoribonuclease NrnB among *Priestia megaterium* and its relatives.** CSI-I and CSI-II indicated two conserved signature indels (CSI) previously mentioned by Gupta *et al.* Boxes with different colours tell the amino acid sequence patterns. The *P. megaterium* type strain was underlined. Red asterisks show the type and assigned NCBI representative strains of *P. aryabhattai*. All other strains can be found in Table S1.
