## Supplementary material for "A cyclic dipeptide for salinity stress alleviation and the trophic flexibility of an endophyte reveal niches in salt marsh plant-microbe interactions": priest01_sup_table.BioRxiv.pdf

**Article title:**

**Affiliations:**

**Table S1. List of the genome sequences used in this study.** Species name abbreviations: *Pm*, *Priestia megaterium*; *Pa*, *Priestia aryabhattai*; *Pf*, *Priestia flexa.* Superscripts following the strain names: T, type strain; R, NCBI representative genome. Category abbreviations: P, plant-associated; I, industrial; C, clinical; E, environmental; O, others. NA indicate the data are missing or inaccessible.

| Species | Strain | Accession | Assembly | Size (Mb) | GC (%) | Geographic origin | Isolation source | Category |
|---|---|---|---|---|---|---|---|---|
| *Pm* | BP01R2 | GCA_022537925 | Complete | 5.61 | 37.50 | Taichung, Taiwan | Endosphere of *Bolboschoenus planiculmis* | P |
| *Pm* | ATCC 14581$^{T}$ | GCA_017086525 | Complete | 5.75 | 37.85 | NA | NA | O |
| *Pm* | S2 | GCA_012275205 | Complete | 6.47 | 38.60 | Russia | NA | I |
| *Pm* | CDC 2008724142 | GCA_017086565 | Complete | 6.00 | 37.78 | Rhode Island, USA | *Homo sapiens* | C |
| *Pm* | BHS1 | GCA_015582655 | Complete | 5.37 | 37.80 | Giresun city, Turkey | Alkaline water | E |
| *Pm* | 2020WEIHUA_L | GCA_022023815 | Complete | 5.92 | 37.56 | China | Soil | P |
| *Pm* | S188 | GCA_011058275 | Complete | 5.41 | 37.91 | Cheonan-si, South Korea | Soil | E |
| *Pm* | FDU301 | GCA_013146705 | Complete | 6.87 | 36.89 | Shanghai, China | Paper surface | E |
| *Pm* | Mes11 | GCA_013458535 | Complete | 6.17 | 37.55 | Limagne, France | Agricultural soil growing maize | P |
| *Pm* | BIM B-1314D | GCA_013389435 | Complete | 5.98 | 37.71 | Salihorsk District, Belarus | Potash salt dump | E |
| *Pm* | 5-3 | GCA_009911775 | Complete | 5.17 | 38.30 | Shangluo, China | Farmland | P |
| *Pm* | KNU-01 | GCA_006385935 | Complete | 5.71 | 37.88 | South Korea | Soil | E |
| *Pm* | IGA-FME-1 | GCA_015643545 | Complete | 5.15 | 38.20 | Lishu, Jilin, China | Bulk soil of maize | P |
| *Pm* | H2 | GCA_017352315 | Complete | 6.41 | 37.49 | Turkey: Tuz Golu | Hypersaline environment | E |
| *Pm* | YC4-R4 | GCA_003072605 | Complete | 5.43 | 38.10 | Fujian, China | Rhizosphere of *Spartina anglica* (Hubb.) grow in salty-soil | P |
| *Pm* | JX285 | GCA_002009195 | Complete | 5.61 | 37.90 | Nanchang, China | Rhizosphere *Camellia oleifera* | P |
| *Pm* | HGS7 | GCA_017798265 | Complete | 5.27 | 38.14 | Chongqing, China | Mulberry | P |
| *Pm* | A | GCA_009497655 | Complete | 5.24 | 38.15 | NA | Plant | P |
| *Pm* | CDC 2008724129 | GCA_017086545 | Complete | 5.61 | 38.08 | Rhode Island, USA | *Homo sapiens* | C |
| *Pm* | NCT-2 | GCA_000334875 | Complete | 5.88 | 37.80 | China | Secondary salinization soil of greenhouse | E |
| *Pm* | DSM 319 | GCA_000025805 | Complete | 5.10 | 38.10 | NA | NA | I |
| *Pm* | Q3 | GCA_001050455 | Complete | 5.23 | 38.27 | Hunan, China | Endosphere of tobacco grown in quinclorac contaminated soil | P |
| *Pm* | QM B1551 | GCA_000025825 | Complete | 5.52 | 37.97 | NA | NA | I |
| *Pm* | WSH-002 | GCA_000225265 | Complete | 5.08 | 38.15 | NA | NA | I |
| *Pm* | 22-2$^{R}$ | GCA_009935415 | Scaffold | 5.60 | 37.60 | China | Commercial probiotic | I |
| *Pa* | B8W22$^{T}$ | GCA_000956595 | Contig | 5.10 | 38.00 | Hyderabad, India | NA | E |
| *Pa* | K13$^{R}$ | GCA_002688605 | Complete | 5.25 | 38.14 | Iksan, South Korea | Compost | I |
| *Pf* | NBRC 15715$^{T}$ | GCA_001591565 | Contig | 3.91 | 37.60 | NA | NA | O |

**Table S2. Co-up-DEGs GO ontologies enrichment.**

| Cluster | GO | Description | *p*-value |
|---|---|---|---|
| Biological Process | 0009664 | plant-type cell wall organization | 4.9E-07 |
| | 0071555 | cell wall organization | 1.8E-06 |
| | 0045229 | external encapsulating structure organization | 7.2E-06 |
| | 0071669 | plant-type cell wall organization or biogenesis | 1.6E-04 |
| | 0071554 | cell wall organization or biogenesis | 2.2E-06 |
| Molecular Function | 0005199 | structural constituent of cell wall | 1.5E-08 |
| | 0016762 | xyloglucan:xyloglucosyl transferase activity | 3.2E-02 |
| | 0004601 | peroxidase activity | 1.0E-02 |
| | 0016684 | oxidoreductase activity, acting on peroxide as acceptor | 1.3E-02 |
| | 0016209 | antioxidant activity | 3.4E-02 |
| | 0046906 | tetrapyrrole binding | 3.3E-03 |
| | 0020037 | heme binding | 3.0E-02 |
| Cellular Component | 0005615 | extracellular space | 3.1E-02 |
| | 0005618 | cell wall | 5.9E-04 |
| | 0030312 | external encapsulating structure | 8.1E-04 |
| | 0005576 | extracellular region | 1.2E-02 |
| | 0043231 | intracellular membrane-bounded organelle | 1.1E-05 |
| | 0043227 | membrane-bounded organelle | 9.1E-06 |
| | 0005622 | intracellular anatomical structure | 7.9E-08 |
| | 0043229 | intracellular organelle | 5.5E-06 |
| | 0043226 | organelle | 4.7E-06 |

**Table S3. Co-down-DEGs GO ontologies enrichment.**

| Cluster | GO | Description | *p*-value |
|---|---|---|---|
| Biological Process | 1900366 | negative regulation of defense response to insect | 2.5E-02 |
| | 2000068 | regulation of defense response to insect | 4.2E-03 |
| | 2000022 | regulation of jasmonic acid mediated signaling pathway | 1.1E-06 |
| | 0071456 | cellular response to hypoxia | 6.4E-22 |
| | 0036294 | cellular response to decreased oxygen levels | 7.8E-22 |
| | 0071453 | cellular response to oxygen levels | 8.5E-22 |
| | 0001666 | response to hypoxia | 9.0E-24 |
| | 0036293 | response to decreased oxygen levels | 1.7E-23 |
| | 0070482 | response to oxygen levels | 2.1E-23 |
| | 0010200 | response to chitin | 7.4E-12 |
| | 0009404 | toxin metabolic process | 1.7E-02 |
| | 0009753 | response to jasmonic acid | 3.4E-17 |
| | 0070542 | response to fatty acid | 4.0E-17 |
| | 0010243 | response to organonitrogen compound | 3.7E-13 |
| | 0009751 | response to salicylic acid | 1.6E-06 |
| | 0009611 | response to wounding | 1.4E-14 |
| | 1901698 | response to nitrogen compound | 7.9E-14 |
| | 0014070 | response to organic cyclic compound | 1.8E-09 |
| | 0031347 | regulation of defense response | 1.6E-08 |
| | 0006979 | response to oxidative stress | 4.3E-06 |
| | 0019748 | secondary metabolic process | 2.9E-09 |
| | 0009414 | response to water deprivation | 2.2E-11 |
| | 0009620 | response to fungus | 1.5E-10 |
| | 0009415 | response to water | 2.4E-11 |
| | 0033554 | cellular response to stress | 6.4E-12 |
| | 0001101 | response to acid chemical | 5.2E-11 |
| | 0080134 | regulation of response to stress | 6.5E-08 |
| | 0006970 | response to osmotic stress | 5.0E-07 |
| | 0009651 | response to salt stress | 1.0E-03 |
| | 0050832 | defense response to fungus | 1.3E-03 |
| | 0070887 | cellular response to chemical stimulus | 1.3E-15 |
| | 0033993 | response to lipid | 7.0E-13 |
| | 1901700 | response to oxygen-containing compound | 7.3E-24 |
| | 0097305 | response to alcohol | 1.5E-06 |
| | 0010035 | response to inorganic substance | 5.4E-14 |
| | 0043207 | response to external biotic stimulus | 4.0E-13 |
| | 0051707 | response to other organism | 4.0E-13 |
| | 0009607 | response to biotic stimulus | 4.2E-13 |
| | 0044419 | biological process involved in interspecies interaction between organisms | 5.3E-13 |
| | 0048583 | regulation of response to stimulus | 4.1E-05 |
| | 0009737 | response to abscisic acid | 1.5E-03 |
| | 0009605 | response to external stimulus | 3.8E-14 |

| | | | |
|---|---|---|---|
| | 0009725 | response to hormone | 6.8E-11 |
| | 0009719 | response to endogenous stimulus | 1.1E-10 |
| | 0042221 | response to chemical | 2.2E-28 |
| | 1901701 | cellular response to oxygen-containing compound | 4.5E-03 |
| | 0010033 | response to organic substance | 2.3E-15 |
| | 0006950 | response to stress | 2.2E-26 |
| | 0098542 | defense response to other organism | 8.0E-07 |
| | 0009628 | response to abiotic stimulus | 1.9E-17 |
| | 0006952 | defense response | 6.8E-08 |
| | 0009617 | response to bacterium | 1.8E-02 |
| | 0071310 | cellular response to organic substance | 4.5E-03 |
| | 0051716 | cellular response to stimulus | 4.1E-12 |
| | 0006082 | organic acid metabolic process | 4.3E-04 |
| | 0043436 | oxoacid metabolic process | 3.5E-03 |
| | 0007154 | cell communication | 5.4E-04 |
| | 0007165 | signal transduction | 1.0E-02 |
| | 0023052 | signaling | 1.5E-02 |
| | 0050896 | response to stimulus | 2.6E-26 |
| | 0044281 | small molecule metabolic process | 1.6E-02 |
| Molecular Function | 0120091 | jasmonic acid hydrolase | 9.3E-03 |
| | 0016491 | oxidoreductase activity | 2.2E-02 |
| Cellular Component | 0032991 | protein-containing complex | 4.1E-02 |

**Table S7. Genes associated with rhizosphere competence, plant growth-promoting and associated, salinity stress alleviation as well and oxygen and carbon resource limitation identified in *Priestia megaterium* BP01R2.**

| Category | Function | Gene name | Description (NCBI Protein, GenBank) | BP01R2 locus_tag (MGJ28_RS) |
|---|---|---|---|---|
| Rhizosphere competence | phosphate solubilization and mineralization | *gcd* | glucose dehydrogenase, PQQ-dependent | 20270 |
| | | *phoP* | two-component response regulator PhoP | 24715, 11185, 15795, 24765 |
| | | *phoR* | two-component sensor histidine kinase PhoR | 10820, 24760, 15790 |
| | | *phoA, phoB* | alkaline phosphatase | 06180 |
| | | *phoD* | alkaline phosphatase D | 26445 |
| | | *ppx* | exopolyphosphatase Ppx | 06650 |
| | nitrogen assimilation and reduction | *narK, nasA* | nitrite extrusion protein | 03965 |
| | | *nasC* | nitrate reductase, catalytic subunit | 03960 |
| | | *nasBD, nirB* | nitrate reductase, electron transfer subunit | 03955, 05855 |
| | | *nasE, nirD* | nitrite reductase [NAD(P)H], small subunit | 05860 |
| | siderophore synthesis/Fe-uptake | *na* | siderophore biosynthesis protein | 21240, 21245, 21250, 21255, 21260, 21265 |
| | | *na* | polyketide synthase | 07325 |
| | | *yfmC* | iron(III)-citrate import ABC transporter, iron(III)-citrate-binding protein | 04445 |
| | | *yfmD* | iron(III)-citrate import ABC transporter, permease protein | 04450 |
| | | *yfmE* | iron(III)-citrate import ABC transporter, permease protein | 04455 |
| Plant-associated | L-tryptophane, indole synthesis | *trpA* | tryptophan synthase, alpha subunit | 22520 |
| | | *trpB* | tryptophan synthase, beta subunit | 22525 |
| | | *trpC* | indole-3-glycerol-phosphate synthase | 22535 |
| | | *trpD* | anthranilate phosphoribosyltransferase | 22540, 16125 |
| | | *trpE* | anthranilate synthase component I | 00470, 22545 |
| | | *trpF* | N-(5'-phosphoribosyl) anthranilate isomerase | 22530 |
| | auxin (indole-3-acetic acid) synthesis | *gatA, iaaH* | glutamyl-tRNA(Gln) and/or aspartyl-tRNA(Asn) amidotransferase, A subunit | 01505, 05255, 17275 |

| | | | | |
|---|---|---|---|---|
| | auxin transport | *na* | auxin efflux carrier (AEC) family transporter | 09980, 14190, 14240, 19315 |
| | acetoin, butanediol synthesis | *alsD, budA, aldC* | alpha-acetolactate decarboxylase | 04010 |
| | | *alsS* | acetolactate synthase, catabolic | 04015, 08580 |
| | | *ilvH, ilvN* | acetolactate synthase, small subunit | 24430 |
| | | *ilvB, ilvG, ilvI* | acetolactate synthase, large subunit, biosynthetic type | 24435, 04830 |
| | | *bdhA* | 2,3-butanediol dehydrogenase | 12195, 25665, 11290, 15390, 11965, 09800 |
| | flagellar assembly | *flgB* | flagellar basal-body rod protein FlgB | 21900 |
| | | *flgC* | flagellar basal-body rod protein FlgC | 21895 |
| | | *flgD* | flagellar hook assembly protein | 21850 |
| | | *flgE* | flagellar hook protein FlgE | 21845 |
| | | *flgK* | flagellar hook-associated protein FlgK | 26530, 10685 |
| | | *flgL* | flagellar hook-associated protein FlgL | 26525 |
| | | *flgM* | negative regulator of flagellin synthesis (anti-sigma-D factor) | 26540 |
| | | *flgN* | conserved hypothetical protein | 26535 |
| | | *flhA* | flagellar biosynthesis protein FlhA | 21790 |
| | | *flhB* | flagellar biosynthetic protein FlhB | 21795 |
| | | *hag, fliC* | flagellin | 05595 |
| | | *fliD* | flagellar hook-associated protein FliD | 26480 |
| | | *fliE* | flagellar hook-basal body complex protein FliE | 21890 |
| | | *fliF* | flagellar M-ring protein FliF | 21885 |
| | | *fliG* | flagellar motor switch protein FliG | 21880 |
| | | *fliH* | flagellar assembly protein FliH | 21875 |
| | | *fliI* | flagellum-specific ATP synthase | 21870 |
| | | *fliJ* | flagellar export protein FliJ | 21865 |
| | | *fliK* | flagellar hook-length control protein | 21855 |
| | | *fliM* | flagellar motor switch protein FliM | 21830 |
| | | *fliN, fliY* | flagellar motor switch protein FliN | 21825 |
| | | *fliP* | flagellar biosynthetic protein FliP | 21810 |
| | | *fliQ* | flagellar biosynthetic protein FliQ | 21805 |
| | | *fliR* | flagellar biosynthetic protein FliR | 21800 |

| | | | | |
|---|---|---|---|---|
| | | *fliT* | flagellar assembly protein FliT | 26470 |
| | | *fliS* | flagellar protein FliS | 26475, 26570 |
| | | *motA* | chemotaxis protein MotA | 10425 |
| | | *motB* | chemotaxis protein MotB | 10430 |
| | bacterial chemotaxis | *cheA* | chemotactic two-component sensor histidine kinase | 21770 |
| | | *cheB* | chemotactic two-component response regulator-glutamate methylesterase | 21775 |
| | | *cheD* | chemoreceptor glutamine deamidase CheD | 21760 |
| | | *cheR* | chemotaxis protein methyltransferase | 07775, 22560 |
| | | *cheY* | chemotaxis protein CheY | 04760, 21820 |
| | | *cheW* | chemotactic signal transduction protein | 21765 |
| Salinity stress alleviation | superoxide dismutase | *sodACF* | superoxide dismutase | 10920, 23510, 25700, 14750 |
| | catalase | *cat* | catalase | 16360, 27180, 20625, 17200, 17205 |
| | spermidine synthesis | *speA* | arginine decarboxylase | 06820, 00240 |
| | | *speB* | agmatinase | 11530, 26925, 04410 |
| | | *speH, speD, AMD1* | S-adenosylmethionine decarboxylase | 24730, 15765 |
| | | *speE, SRM* | spermidine synthase | 26930, 03970 |
| Oxygen and carbon resource limitation | anaerobic respiration | *ldh* | L-lactate dehydrogenase | 02650 |
| | | *alsS* | acetolactate synthase AlsS | 04015 |
| | | *budA* | acetolactate decarboxylase | 04010 |
| | | *pta* | phosphate acetyltransferase | 26985 |
| | | *na* | acetate kinase | 24900 |
| | | *ar* | 2,3-butanediol dehydrogenase | 09800 |
| | | *acoABC* | acetoin dehydrogenas | 09775, 09780, 09785 |
| | | *pdh* | pyruvate dehydrogenase | 06785, 06790, 06795, 03690, 06800 |
| | | *acsA* | acetyl-CoA synthetas | 12695, 24990 |
| | anaerobic regulatory | *resA* | thiol-disulfide oxidoreductase ResA | 22785 |
| | | *na* | cytochrome c biogenesis protein ResB | 22780 |

| | | | | |
|---|---|---|---|---|
| | | *ccsB* | c-type cytochrome biogenesis protein CcsB | 22775 |
| | | *na* | response regulator transcription factor | 22770 |
| | | *na* | ATP-binding protein | 22765 |
| | | *fnr* | Fnr family transcriptional regulator | 03420 |
| | reductive glycine (rGly) pathway | *fdhF* | formate dehydrogenase subunit alpha | 12430, 12505, 26155 |
| | | *fdhD* | formate dehydrogenase accessory sulfurtransferase FdhD | 26145, 12455 |
| | | *focA* | formate transporter | 05870, 12745 |
| | | *ftl* | formate-tetrahydrofolate (THF) ligase | 10120 |
| | | *folD* | methenyltetrahydrofolate cyclohydrolase/methylenetetrahydrofolate dehydrogenase (NADP+) | 10735 |
| | | *gcvPA* | glycine dehydrogenase (glycine cleavage system P1 protein) | 23360 |
| | | *gcvPB* | glycine dehydrogenase (glycine cleavage system P2 protein) | 23355 |
| | | *gcvH* | glycine cleavage system protein H-protein | 25960 |
| | | *gcvT* | glycine cleavage system T-protein | 23365 |
| | | *shmt* | serine hydroxymethyltransferase | 26770 |
| | | *sdaAB* | L-serine ammonia-lyase, iron-sulfur-dependent subunit beta | 22055 |
| | | *sdaAA* | L-serine ammonia-lyase, iron-sulfur-dependent, subunit alpha | 22050 |
| | | *na* | D-serine ammonia-lyase | 10635 |
| | reductive tricarboxylic acid (rTCA) pathway | *pyc* | pyruvate carboxylase | 06900 |
| | | *ppc* | phosphoenolpyruvate carboxylase | 04135 |
| | | *mdh* | malate dehydrogenase | 24775 |
| | | *fum* | fumarate hydratase | 01970, 11865 |
| | | *sdhABC* | succinate dehydrogenase/fumarate reductase | 24550, 24555, 24560 |
| | | *sucCD* | succinate-CoA ligase | 21940, 21945 |
| | | *korAB* | 2-oxoglutarate ferredoxin oxidoreductase | 21490, 21485 |
| | | *icd* | NADP-dependent isocitrate dehydrogenase | 24780 |
| | | *acnA* | aconitate hydratase AcnA | 13820 |
| | | *cs* | citrate synthase CS | 16565, 24785 |

| | | | | |
|---|---|---|---|---|
| | ferredoxin | *fer* | ferredoxin | 12115, 17790, 22745, 04525, 12240 |
